# The *E. coli* PeptideAtlas Build: Characterizing the observed *Escherichia coli* pan-proteome and its post-translational modifications

**DOI:** 10.1101/2025.09.11.675345

**Authors:** Caroline Jachmann, Zhi Sun, Kevin Velghe, Florence Arsène-Ploetze, Aurélie Hirschler, Jasper Zuallaert, Christine Carapito, Robbin Bouwmeester, Kay Nieselt, Eric Deutsch, Lennart Martens, Ralf Gabriels, Tim Van Den Bossche

## Abstract

*Escherichia coli* is a widely used model organism in molecular biology. Despite its pivotal role, a comprehensive proteome resource covering the *E. coli* pan-proteome and its post-translational modifications (PTMs) has been lacking. Here we present the *E. coli* PeptideAtlas build, the first comprehensive pan-proteome analysis of *E. coli,* generated from 40 high-quality public and in-house datasets spanning a broad diversity of strains, sample types, and experimental conditions, and comprising over 73 million MS/MS spectra. All datasets were reprocessed using both a closed search (Trans-Proteomic Pipeline using MSFragger) and an open search (ionbot). The *E. coli* PeptideAtlas build provides evidence for 4,755 proteins, including 1,410 previously lacking protein-level support in UniProt. The resource offers protein coverage, modification sites, raw spectra with matched peptides, and manually annotated metadata for the *E. coli* pan proteome. PTM profiling identified over 10,000 modification sites, including phosphorylation (3,806), acetylation (754), methylation (730), glutathionylation (352) and phosphoribosylation (226). Analysis of the glutathionylation sites revealed potential links to metal binding regulation. We also detected proteins likely associated with phages, underscoring the value of pan-proteomic approaches for studying host-phage interactions. All identifications are publicly accessible and traceable through the PeptideAtlas interface. We expect that the *E. coli* PeptideAtlas build will provide a useful resource for the community, which supports, for example, targeted MS experiment design, PTM enrichment method development, and strain typing. It allows straightforward lookups of protein and peptide identifications and facilitates comparative proteomic analyses by enabling the assessment of protein presence and variability across different *E. coli* strains. The build is available at https://db.systemsbiology.net/sbeams/cgi/PeptideAtlas/buildDetails?atlas_build_id=585.

## Introduction

*Escherichia coli* (*E. coli*) is a cornerstone in biological research, serving as a key model organism in various fields, including biotechnology and biopharmaceutical research^1^. Its extensive biological diversity ranges from probiotic strains beneficial to humans to pathogenic strains associated with diseases such as diarrhea, urinary tract infections, and meningitis^2^. The species exhibits vast genomic diversity, with a pan-genome encompassing up to 18,000 genes^3^, of which only a small portion is shared across all strains. This genetic variability is driven by the species’ high genomic plasticity^1^, facilitating rapid adaptation via horizontal gene transfer.

Despite the wealth of genomic information, proteomics data for *E. coli* remain fragmented. Most studies focus on a few K-12 derivatives, using the MG1655 reference proteome. The largest *E. coli* proteome profiling studies to date have identified up to 3,300 proteins^4^, with quantification data available for 2,300 proteins^5^. These studies have examined the *E. coli* proteome under diverse conditions, including different growth media and stages, which is essential for detecting condition-specific proteins. Smaller scale projects have also explored specific conditions like antibiotic stress^6^, high temperature^7^, and even spaceflight^8^.

Thanks to the adoption of open data standards in proteomics, these and many other *E. coli* datasets are now available in public repositories like ProteomeXchange. While these data are valuable, they often lack reprocessing or integration into comprehensive resources. Indeed, while *E. coli* is the most represented prokaryote in ProteomeXchange, no unified reprocessing has been conducted until now. Moreover, existing resources such as PeptideAtlas provide valuable tools for building proteome profiles, but have yet to offer a comprehensive *E. coli* resource, especially one that captures the protein and post-translational modification (PTM) landscape in multiple strains and conditions. And perhaps most importantly, many protein entries in UniProt still lack experimental evidence at the protein level, limiting their utility in functional or comparative studies.

In this study, we present the first systematic reprocessing effort to characterize the expressed *E. coli* pan-proteome and its modifications. By integrating public and in-house LC-MS data with the Trans-Proteomics Pipeline^9^ (TPP) and ionbot^10^, an open-modification search engine, we created the *E. coli* PeptideAtlas build. This resource includes both core and strain-specific proteins, providing MS-based evidence for protein entries in UniProt previously annotated as ‘predicted’ or lacking experimental validation. We also identified a wide range of PTM types and modification sites, contributing to their functional annotation. The PeptideAtlas build is publicly accessible and supports various applications, including targeted MS assay development, method development for PTM enrichment, and comparative proteomic analyses across different *E. coli* strains.

## Methods

### Data collection

LC-MS/MS projects available through ProteomeXchange were filtered for 1) only containing *E. coli*, and 2) data acquisition performed on Orbitrap instruments (Q Exactive, LTQ Orbitrap), with the exception of PXD020785, a large spectral library assay acquired on TripleTOF instruments. The resulting projects were then manually filtered for projects using HCD fragmentation and trypsin or a combination of trypsin and LysC as a cleavage agent, ensuring compatibility with ionbot (again, with the exception of PXD020785, which was only analysed with TPP). Metaproteomics, crosslinking, and incorrectly labelled experiments were excluded. These publically available data were supplemented with in-house acquired data for *E. coli* strain W3110^11^ under six different conditions (see below for details).

Table 1 shows an overview of the 40 included projects, a more extensive description including e.g. utilized growth media and stages available in Supplementary Table 1. Employed methodologies include solubility-based fractionation (1 project), pH-based fractionation (2 projects), and pre-enrichment for certain modifications and cellular components, such as phosphorylated peptides, lactyllysine-modified peptides, outer membrane proteins, and membrane vesicles.

**Table 1:** Included projects with their corresponding treatments and enrichment strategies.

| Project ID | Strain / isolate | Isolate from | Patho genic | Treatment | Enrichment | Labeling | Ref |
| --- | --- | --- | --- | --- | --- | --- | --- |
| PXD058808 | Escherichia coli str. K-12 substr. W3110 |  | no | Different growth media |  |  | this paper |
| PXD000498 | Escherichia coli str. K-12 substr. MG1655, Escherichia coli str. K-12 substr. NCM3722, Escherichia coli BW25113 |  | no | Temperature stress |  |  | ref <sup>12</sup> |
| PXD007864 | Escherichia coli str. K-12 substr. MC4100 |  | no |  | CobB Affine proteins |  | ref <sup>13</sup> |
| PXD008369 | Escherichia coli str. K-12 substr. W3110 |  | no |  | Phosphorylation |  | ref <sup>14</sup> |
| PXD008921 | Escherichia coli str. K-12 substr. W3110 |  | no | Antibiotic stress | Phosphorylation |  | ref <sup>15</sup> |
| PXD011023 | Escherichia coli str. K-12 substr. MG1655 |  | no | Pressure |  |  | ref <sup>14</sup> |
| PXD012611 | Escherichia coli str. K-12 substr. MG1655 |  | no |  |  |  | ref <sup>7</sup> |
| PXD013273 | Escherichia coli str. K-12 substr. MG1655 |  | no | Different detergents |  |  | ref <sup>16</sup> |
| PXD013610 | Escherichia coli BW35113 |  | no |  |  |  | ref <sup>17</sup> |
| PXD016001 | Escherichia coli str. W0153 |  | no | Antibiotic stress |  |  | ref <sup>6</sup> |
| PXD016074 | Escherichia coli str. W0153 |  | no | Antibiotic stress |  |  | ref <sup>6</sup> |
| PXD016669 | Escherichia coli DH5 |  | no |  |  |  | ref <sup>18</sup> |
| PXD017618 | Escherichia coli str. K-12 substr. MG1655 |  | no |  |  |  | ref <sup>19</sup> |
| PXD019140 | Escherichia coli str. K-12 |  | no |  |  |  | ref <sup>20</sup> |
| PXD019288 | Not available |  | no | Copper stress |  |  | ref <sup>21</sup> |
| PXD020207 | Escherichia coli str. O2:K1; ST95, Escherichia coli str. O6:K2; ST73, Escherichia coli str. O2:K1; ST117 | chicken, human blood | yes, ExPEC |  | Extracellular Vesicles |  | ref <sup>22</sup> |
| PXD020249 | Escherichia coli str. K-12 substr. MG1655 |  | no | auxotroph, anaerobic, aerobic conditions | Nitrosylation | SILAC | ref <sup>23</sup> |
| PXD020785 | ASKA(-) library Host Cell K-12 AG1(ME5305), Escherichia coli str. K-12 |  | no |  |  |  | ref <sup>4</sup> |
| PXD022070 | Escherichia coli BW25993 |  | no |  |  |  | ref <sup>24</sup> |
| PXD022526 | Escherichia coli str. K-12 substr. MG1655 |  | no |  |  |  | ref <sup>25</sup> |
| PXD023158 | Escherichia coli str. K-12 |  | no | Spaceflight |  | TMT | ref <sup>8</sup> |
| PXD024138 | Escherichia coli ATCC 25922 |  | no | Plant Defensin stress |  |  | ref <sup>26</sup> |
| PXD025088 | Escherichia coli NC101 | mouse | yes, AIEC | Antibiotic stress |  |  | ref <sup>27</sup> |
| PXD025706 | Not available |  | no | Antioxidant stress |  |  | ref <sup>28</sup> |
| PXD029347 | APEC strain AE17(O2 serotype) | chicken | yes, APEC |  | Membrane Vesicles |  | ref <sup>29</sup> |
| PXD029931 | Escherichia coli str. K-12 substr. MG1655 |  | no | Removal of oriC |  |  | ref <sup>30</sup> |
| PXD030345 | Escherichia coli str. K-12 substr. MG1655 |  | no | cobB knockout | Lactyllysine | SILAC | ref <sup>31</sup> |
| PXD030346 | Escherichia coli str. K-12 substr. MG1655 |  | no | yiaC knockout | Lactyllysine | SILAC | ref <sup>31</sup> |
| PXD030650 | E. coli BA102 |  | no | Gene addition |  | TMT | ref <sup>32</sup> |
| PXD030690 | Escherichia coli str. K-12 |  | no | Toxin stress |  | iTRAQ | ref <sup>33</sup> |
| PXD031694 | Escherichia coli BL21(DE3) |  | no |  |  |  | ref <sup>34</sup> |
| PXD032954 | Not available |  | no |  |  |  | ref <sup>35</sup> |
| PXD035590 | Not available |  | no |  |  |  | ref <sup>36</sup> |
| PXD036749 | Escherichia coli str. JW2806, Escherichia coli str. K-12 substr. W3110 |  | no |  | DDM-and-Colicin-Streptavidin-Pull down, Triton-then-LDA O | SILAC | ref <sup>37</sup> |
| PXD037221 | Escherichia coli str. K-12 substr. W3110 |  | no |  |  | TMT | ref <sup>38</sup> |
| PXD040621 | Escherichia coli O127:H6 strain E2348/69 |  | yes, EPEC | Antioxidant stress |  |  | ref <sup>39</sup> |
| PXD041301 | Escherichia coli str. DSM 105380 | blood from unknown organism | yes |  | Membrane Vesicles |  | ref <sup>40</sup> |
| **PXD043473** | Escherichia coli BL21(DE3) |  | no |  |  |  | ref <sup>41</sup> |
| **PXD045084** | Escherichia coli str. K-12 substr. MG1655 |  | no | Toxin stress |  |  | ref <sup>42</sup> |
| **PXD050358** | Escherichia coli ST93, Escherichia coli O25b:H4-ST131, Escherichia coli ST73, Escherichia coli ATCC 25922, Escherichia coli ST1421 | clinical isolates | yes | Antibiotic stress |  |  | ref <sup>43</sup> |

### LC-MS/MS acquisition

In addition to public data, we also included in-house data acquired on different growth media and growth stages. First, 50 ml of M9 + glucose (2 g/L) were inoculated from a 5 ml culture of E. coli strain W3110 grown overnight in LB and incubated at 37°C with agitation. On the second day, this 50 ml culture was used to inoculate two different 100 ml media (LB (LB Lennox Agar Broth, Euromedex, Souffelweyersheim, France) or M9 (M9CA Medium, Euromedex, Souffelweyersheim, France) + glucose (2 g/l)) to obtain an initial OD of 0.01 (in LB) or 0.02 (M9 + glucose), with four replicates each. The cells were cultured at 37°C with agitation. After 8 hours, the 100 ml cultures of M9 + glucose reached an OD of approximately 0.1. 2 x 50 ml of each of these cultures were centrifuged for 10 minutes at 5000 rpm. One of the two pellets was resuspended in 50 ml of M9 + glucose (2 g/L) and the other pellet was resuspended in 50 ml + acetate (2 g/L) and incubated at 37 °C with agitation. Under each culture condition, cells were sampled at two growth stages (exponential growth stage, OD ≈ 0.6, and late stationary phase, 72 h).

Proteins were extracted from frozen cell pellets by resuspension in Laemmli like buffer, sonication with Bioruptor Pico (10 times, 30s on/off) and centrifugation (5 min, 10,000 g). Protein concentration was measured using the Pierce 660nm Protein Assay kit. In-gel protein digestion was performed according to standard protocols. Briefly, the samples were heated for 5 min at 95 °C, loaded on in-house stacking gels (40 μg), and the gels were run at 40V until proteins were migrated around 1 cm into the gel. Gels were fixated for 15 min, stained with Silver Blue for 1 h, and the bands were excised and transferred to 96-well plate. Proteins were decolored 4x with ACN/NH4HCO3, dehydrated with 100 μl ACN for 5 minutes, reduced with 60μl of 10 mM DTT for 30 min at 60 °C and 30 min at room temperature. Proteins were alkylated with 60 μl of 55mM IAA for 20 min in the dark, washed 4x with 80 μl NH4HCO3, 80μl of ACN for 5 min each, and dried with 80 μl of ACN twice for 5 min. Then, proteins were digested overnight at 37°C using modified porcine trypsin with a final trypsin/protein ratio of 1/100 (Promega, Madison, USA). Peptides were extracted by adding 100 μl ACN (60%) for 1 h under gentle shaking, and 60 μl ACN (100%) for 10 min without. Peptides were dried in a vacuum concentrator and dissolved in 100 μl H20, 0.1% FA, 2% ACN.

Data-dependent acquisition was done on a nanoAcquity Ultra Performance LC device (Waters Corporation, Milford, MA) coupled to a quadrupole-Orbitrap mass spectrometer (Q-Exactive HF-X, Thermo Fisher Scientific, Waltham, MA). A 58 min stepwise gradient was applied (0.1% FA in water (solvent A) and 0.1% FA in ACN (solvent B), 1-35%B). MS1 spectra were acquired at a resolution of 60,000, MS2 spectra at 15,000. Peptide fragmentation was performed using HCD (NCE: 27%). Dynamic exclusion time was set to 30s. The metadata was annotated with lesSDRF^44^. Raw files and associated SDRF^45^-formatted metadata were uploaded to PRIDE and are available under the identifier PXD058808.

### Metadata collection and annotation

Information about all publically available samples and experiments were also annotated in SDRF format with lesSDRF. These machine readable files include information necessary for reprocessing (mass tolerances, labeling, enrichment, among others) and for downstream analyses and interpretation (such as growth media, strain, treatment) down to the level of each MS run. Where necessary, original authors were contacted for more information. These metadata are available in Supplementary Table 1.

### Construction of the proteome search space

To capture sequence variation while limiting the search space and reducing false peptide-spectrum matches, an *E. coli* pan-proteome search database was constructed by combining multiple strain proteomes. Representative strain proteomes were sourced from UniProt whenever available. For strains without publicly available proteomes, closely related strains were identified through literature searches. Seven strains had available proteomes: MG1655, DH5α, BW25113, MC4100, ATCC 25922, BL21, and W3110. For W0153 and BW25993, no direct proteomes were found, but BW25993 was closely related to BW25113 (already included), and for W0153, the proteome of its parent strain AB1157 was included. No proteomes were available for AE17, DSM 105380, BA102, and AG1 (ME5305). For the clinical samples in PXD050358, the whole-genome sequencing-based proteomes provided in the original study were included.

SwissProt bacteriophage proteins were also included to account for phage-derived proteins that may be present in *E. coli* strains. Additionally, known contaminants (excluding *E. coli* proteins) were incorporated from Frankenfield et al.^46^.

The final FASTA search database was constructed by concatenating all selected proteomes. After removing duplicate sequences, this yielded a total of 28,580 target sequences. To ensure robust false discovery rate (FDR) control, an equal number of decoy sequences (28,580) was generated. A complete list of included proteomes, along with their UniProt accession numbers and download dates, is provided in Table 2. The FASTA database itself (with and without decoy protein sequences added) is available for download in the github repository (https://github.com/CompOmics/ecoli-peptideatlas-manuscript/).

**Table 2:** Proteomes included in the search database for the *E. coli* PeptideAtlas build. WGS = Whole genome sequencing.

| Proteome ID | Download date | Organism / *E. coli* strain | # unique proteins | # proteins |
| --- | --- | --- | --- | --- |
| UP000869295 | 13-03-23 | AB1157 | 2944 | 4627 |
| UP000008216 | 08-05-24 | APEC O1:K1 | 2962 | 4870 |
| UP000014099 | 23-02-23 | ATCC25922 | 1298 | 4901 |
| UP000002032 | 07-05-24 | BL21 | 655 | 4156 |
| UP000029103 | 23-02-23 | BW25113 | 4 | 4095 |
| UP001065293 | 07-11-23 | DH5 | 545 | 4223 |
| UP000000625 | 08-02-24 | MG1655 | 102 | 4362 |
| UP000001478 | 02-11-23 | MC4100 | 88 | 4042 |
| UP000005190 | 23-02-23 | NC101 | 2065 | 4728 |
| UP000036496 | 02-11-23 | NCM3722 | 292 | 4504 |
| UP000008205 | 08-05-24 | O127:H6 | 2652 | 4594 |
| UP000868987 | 02-11-23 | W3110 | 193 | 4324 |
| Recacha et al.<sup>43</sup> |  | (WGS, ST73) | 456 | 3703 |
| Recacha et al.<sup>43</sup> |  | (WGS, ST131) | 2266 | 3927 |
| Recacha et al.<sup>43</sup> |  | (WGS, ST1421) | 1232 | 3969 |
| Recacha et al.<sup>43</sup> |  | (WGS, ST93) | 641 | 3943 |
| UP000001156 | 07-02-24 | T1 phage | 6 | 6 |
| UP000503557 | 07-02-24 | T2 phage | 1 | 1 |
| UP000002092 | 07-02-24 | T3 phage | 1 | 1 |
| UP000009087 | 07-02-24 | T4 phage | 268 | 268 |
| UP000502759 | 07-02-24 | T6 phage | 1 | 1 |
| UP000000840 | 07-02-24 | T7 phage | 56 | 57 |
| UP000001711 | 07-02-24 | lambda phage | 13 | 66 |
| Frankenfield et al.<sup>46</sup> |  | Contaminants | 428 | 428 |

### LC-MS/MS data processing

The public and in-house data sets were processed both in a closed search approach (TPP, with MSFragger^47^ as the search engine), and in an open search approach (ionbot^10^).

For TPP reprocessing, the raw mass spectrum files were downloaded from the ProteomeXchange repository and converted into mzML format^48^. The conversion process utilized ThermoRawFileParser^49^ for Thermo files, while SCIEX MS Data Converter 1.3.1^50^ was used for .wiff files from SCIEX instruments. Data analysis was performed using TPP with MSFragger as the search engine, leveraging parameters sourced from the sample data relationship format (SDRF) files associated with each dataset. Where specific parameters were not detailed, default settings were applied. The analysis included a fixed modification of +57.021464 for carbamidomethylated cysteine and variable modifications of +15.9949 for oxidized methionine, +42.0106 for protein N-terminal acetylation, -17.02650 for pyroglutamic acid formation from glutamine and for cyclization of N-terminal S-carbamoylmethyl-cysteine, -18.01060 for pyroglutamic acid formation from glutamic acid, and +0.984016 for deamidation. In cases where experiments involved isobaric or metabolic labeling strategies, labeling-specific modifications were added accordingly. Precursor mass tolerance was set to 10ppm. Fragment mass tolerance was set to 20 ppm, except for PXD030345 and PXD030346, where it was set to 0.6 Da (due to fragment ion analysis in the ion trap). For PXD020785, both precursor and fragment mass tolerances were set to 50ppm. The minimum peptide length was set to seven amino acids. Up to two missed cleavages were allowed, and semitryptic cleavage was used.

Additionally, an open search was performed with ionbot v0.11.4. The raw files were converted to .mgf format using ThermoRawFileParser. Carbamidomethylation of cysteine and oxidation of methionine were set as variable modifications. Additionally, fixed and variable modifications were set accordingly for projects using isobaric labels and PTM enrichment. Semi-tryptic peptides were included, and a maximum number of two missed cleavages were allowed. Precursor and fragment mass tolerances were left at the default values (20 ppm), and rescoring was performed with Percolator^51^. A very strict PSM FDR threshold of 0.05% per run was applied to only include highly confident identifications. Additionally, modifications were relocalised with pyAscore^52,53^, and only PSMs with high confidence (Ascore >20) were integrated into the PeptideAtlas build.

### Creating the PeptideAtlas build

The results from the closed and open search were combined into one PeptideAtlas build available under https://db.systemsbiology.net/sbeams/cgi/PeptideAtlas/buildDetails?atlas_build_id=585. In case of conflicting peptidoform assignments between ionbot and TPP, the closed search result was used. After merging the PSMs, MAYU^54^ was used to ensure a global protein FDR below 1%. PeptideProphet^55^ was employed to evaluate the confidence of PSM identifications, providing probabilities to each identified peptide.

When working with multiple proteomes, PeptideAtlas employs a designated *core proteome* that takes precedence during protein inference. This should not be confused with the core proteome in the context of pan-proteomics, which refers to the set of proteins shared across all strains. For this, we chose UP00000625 (K-12 MG1655), as it is the reference proteome for *E. coli* in UniProt and is assumed to be close to completely annotated^56^.

Depending on their evidence, protein identifications are categorized into one of ten tiers in PeptideAtlas builds. For further analysis in this paper, we use a simplified version of the PeptideAtlas tier system similar to a previous PeptideAtlas build^57^ (Table 3): The “representative” tier was removed as there were no proteins in this tier in this build. Identical proteins are assigned to the tier of their (identical) partner, but are not counted again for summary statistics. Tier 5, 6, and 8 are combined into one Tier (“uncertain”).

**Table 3:**
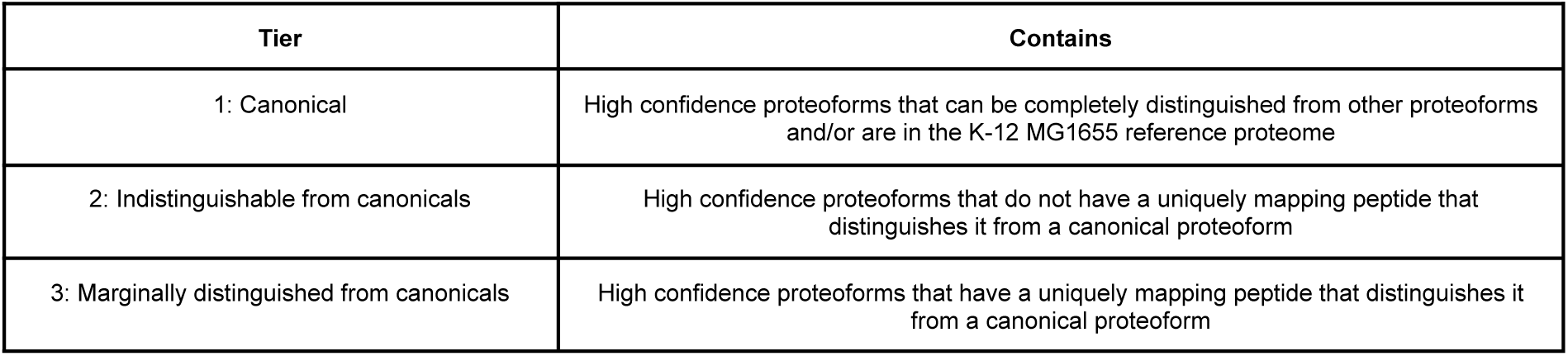
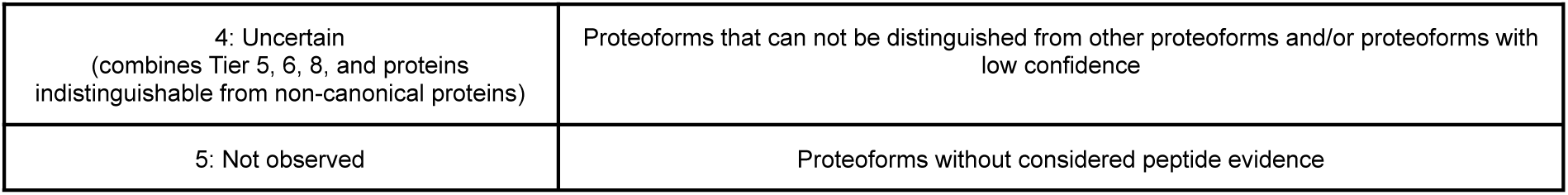
Simplified confidence tiers for this study.

| Tier | Contains |
| --- | --- |
| 1: Canonical | High confidence proteoforms that can be completely distinguished from other proteoforms and/or are in the K-12 MG1655 reference proteome |
| 2: Indistinguishable from canonicals | High confidence proteoforms that do not have a uniquely mapping peptide that distinguishes it from a canonical proteoform |
| 3: Marginally distinguished from canonicals | High confidence proteoforms that have a uniquely mapping peptide that distinguishes it from a canonical proteoform |
| 4: Uncertain<br>(combines Tier 5, 6, 8, and proteins<br>indistinguishable from non-canonical proteins) | Proteoforms that can not be distinguished from other proteoforms and/or proteoforms with<br>low confidence |
| 5: Not observed | Proteoforms without considered peptide evidence |

### Terminology

The search proteome includes multiple strain proteomes and, consequently, contains a substantial proportion of proteins with highly similar sequences. We will therefore refer to FASTA entries as proteoforms in this paper. Clustering of proteoforms gives rise to homology clusters, representing proteoforms of the same protein. There is no universally agreed on similarity threshold to choose for protein clustering, and for this paper we will use 70% as default, as done in Broadbent *et al*.^58^. Homology clusters with at least one proteoform in every included strain are taken as “core proteins”; clusters with at least one proteoform in at least two, but not all strains are referred to as “accessory proteins”, and proteins only found in one strain proteome are called “orphan proteins”.

### Proteome characterization and PTM analysis

Protein subcellular localisations were predicted with PSORTb 3.0^59^, isoelectric points and weights were predicted and calculated, respectively, with Pyteomics^60^. Proteins identified with high confidence (termed canonical proteins) in the build were aligned to all UniProt *E. coli* proteins with known evidence (existence:1 and taxonomy_id:562, 4,351 proteins as of 27.9.24) to evaluate how many proteins gained new experimental evidence. For this, any canonical proteins that did not match with a similarity of 70% or higher to the already observed proteins were considered new. To estimate proteome similarities, pairwise Jaccard similarities were calculated on tryptic digests. For the PTM site analyses, peptides carrying modifications were remapped to canonical proteins to filter out modification sites that are not uniquely mapping to one protein. Protein abundance was estimated by the normalised spectrum abundance factor (NSAF), calculated by counting the number of PSMs mapping uniquely to the protein, and dividing it by the length of the protein. Additionally, the NSAF was normalized by the number of all PSMs found in the run. Batch effects in modification patterns were investigated using t-SNEs, which were calculated based on the observation counts for each PTM per run, i.e. each run is represented by a vector specifying how many times each PTM was found in that run. We employed ESMFold^61^ (esm.pretrained.esmfold_v1 model) for protein structure prediction on account of its computational efficiency.

## Results

### The *E. coli* PeptideAtlas build

To support high-confidence and strain-resolved *Escherichia coli* proteomics, we constructed a comprehensive *E. coli* PeptideAtlas build by integrating publicly available LC-MS/MS datasets from ProteomeXchange with in-house generated data (Figure 1A). The selected 40 projects encompass a broad array of experimental conditions and analytical strategies, including diverse *E. coli* strains, antibiotic and environmental perturbations, sample preparation protocols, and enrichment methods designed to enhance the detection of low-abundance or modified peptides (Figure 1B). In total, over 73 million spectra were reanalyzed using a dual search strategy: a closed search with MSFragger via the Trans-Proteomic Pipeline (TPP) and an open search with ionbot (Figure 1C). These results were then combined under strict protein-level false discovery rate (FDR) control using MAYU.

**Figure 1:**
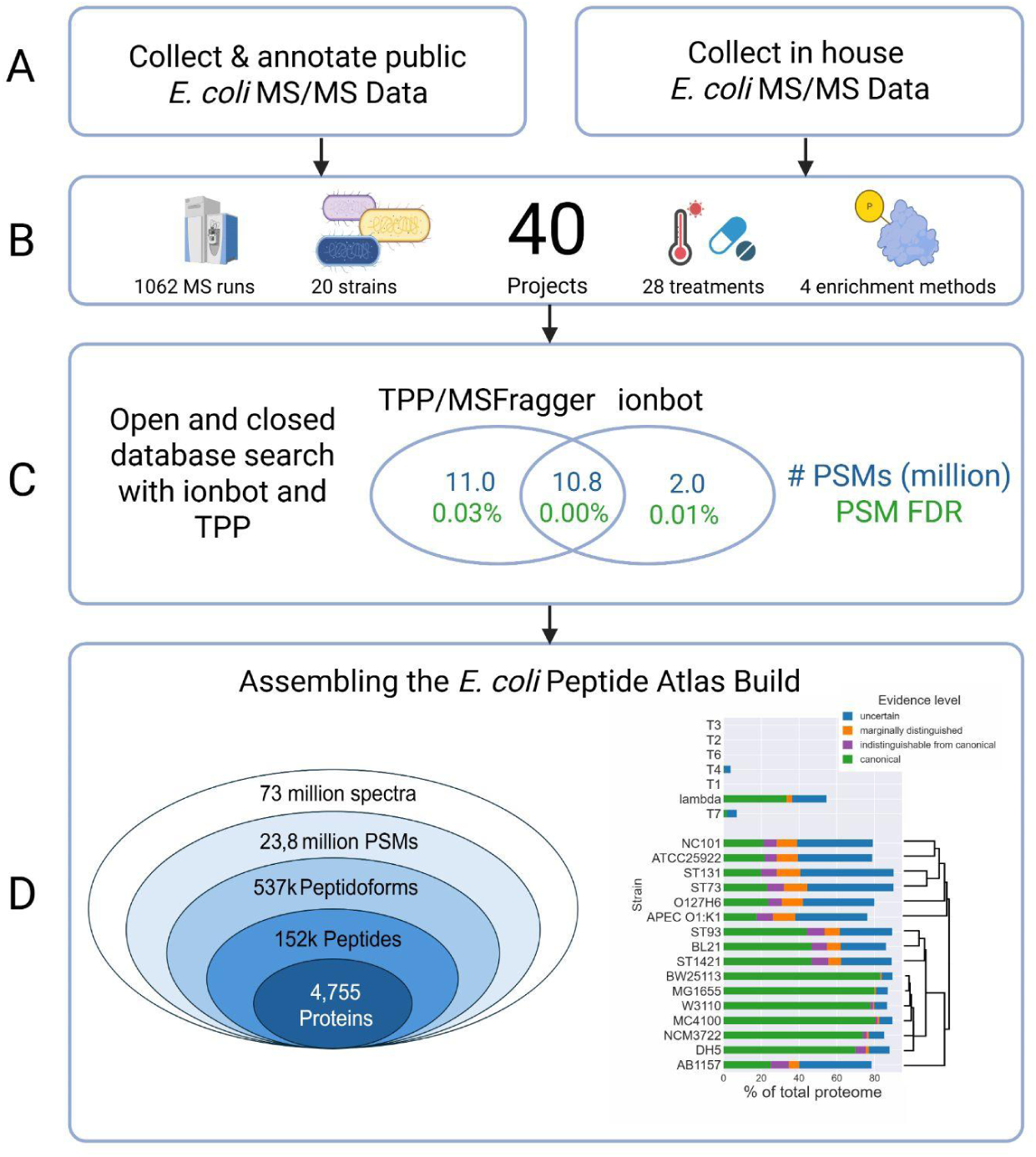
The E. coli PeptideAtlas build. **A:** E. coli PeptideAtlas build created by combining public proteomics data from ProteomeXchange and LC-MS/MS data acquired in-house. **B:** Selected experiments cover a large variety of strains, treatments, sample preparation and enrichment protocols. **C:** Spectra were reprocessed with closed MSFragger search via TPP and open search with ionbot, and results were combined under strict global protein FDR control with MAYU. **D:** 537.885 distinct peptidoforms were identified, giving evidence to 4,755 proteins in the highest confidence tier. Proteome coverages vary between 76-90% across strains including all confidence tiers. For the highest confidence tier, coverages drop for strains not derived from K-12 as K-12 MG1655 proteins were prioritized in case of indistinguishable proteoforms during protein inference. The tree next to the proteome coverage barplot is based on the distance of tryptic digests between the proteomes.

This yielded over 537,000 distinct peptidoforms and provided peptide-level evidence for 4,755 canonical proteins (Figure 1D). The coverage across reference proteomes varies between 76–90%, with a higher canonical coverage in K-12-derived strains due to prioritization of K-12 MG1655 proteins during inference in cases of shared peptide evidence. The hierarchical relationship between proteomes, based on in silico tryptic peptide overlap, underscores both their diversity and the value of incorporating multiple strain backgrounds (Figure 1D, right). This build not only provides a curated, high-resolution view of the *E. coli* proteome but also supports functional annotation and systems-level exploration by linking identified peptides and proteins to experimental metadata and external databases.

#### The selected 40 projects cover a large variety of experiment types

The projects collected from public repositories or generated in house showcase the diverse experimental approaches applied to *E. coli* research globally, encompassing a variety of enrichment and fractionation techniques tailored to enhance the detection of low-abundance peptidoforms and traditionally elusive proteins via LC-MS/MS (see Figure 1B).

These studies investigated proteome changes under numerous environmental conditions, including exposure to 21 different antibiotics and physical conditions like high temperature (42 °C), increased pressure, and spaceflight. Contributions have come from around the globe, with a significant number from Europe (14), followed by Asia (13), America (8), Oceania (2), and Africa (1). Labeling techniques employed in these studies include tandem mass tags (TMT) (4), stable isotope labeling by amino acids in cell culture (SILAC) (3), and isobaric tags for relative and absolute quantitation (iTRAQ) (1).

Both laboratory strains and clinical isolates are represented, with a predominance of *E. coli* K-12 derivatives like MG-1655 (9 projects) and W3110 (5 projects). Additional lab strains used include BLR(DE3), HMS174(DE3), BL21(DE3), and ATCC 25922. Pathogenic groups represented include AIEC, APEC, and ExPEC, with samples sourced from chickens, humans, and mice.

Some studies induced genetic modifications such as gene knockdowns (1), gene knockouts (7), and the introduction of plasmids (5). The proteins encoded by these artificially added genes are not included in this study. Further information on the included plasmids and gene edits are listed in Supplementary Table 1.

To ease exploration of the results, each of the 40 projects were further divided into smaller experiments to account for different experimental setups such as growth media, fractions, strains, and treatments. All experiments are listed in Supplementary Table 2.

#### Combining closed and open search method leads to high identifications at controlled FDR

The curated 40 projects sum to 73,008,556 spectra from 1062 MS runs. TPP and ionbot identified 23,823,673 (33% of all spectra) PSMs at a 1% global protein FDR, mapping to 537,885 distinct peptidoforms and 151,590 distinct peptides (Figure 1C and D). Around half of the PSMs (10.8 million) included in the build were identified in both the open and closed search, and TPP identified an additional 10.8 million PSMs. The open search identified an additional 2 million PSMs from spectra previously unassigned by TPP, 28.6% (576,444) of which contained modifications not considered in the closed search. For the 3,051,960 PSMs with conflicting peptidoform assignments between ionbot and TPP, the peptidoforms assigned by TPP in the closed search were added to the build.

Based on the collected peptide evidence, a total of 4,755 proteins were identified with high confidence. All results are available online through the build page on the PeptideAtlas website, presenting protein overviews with coverages, and strain and experiment provenance information. It is also possible to trace back evidence to original spectra, and to inspect annotated spectra in a viewer. Moreover, cross-references to databases such as UniProt, KEGG^62^, and NCBI^63^ enable users to access additional information on existing biological knowledge, including details on PTMs.

The spectrum matching rates differ between experiments (subsets of runs in a project with the same experimental setup), from lower than 1% of spectra identified in single cell runs, to 80% of spectra acquired in standard bulk setups. The median ID rate is 42.8%, with higher ID rates for runs on Q Exactive instruments, and lower rates on Orbitrap Fusion, Eclipse, and Velos (Supplementary Figure 1). We observed more distinct peptides in experiments where Lys-C was used in combination with trypsin (median 45.480 peptides per experiment) than when only using trypsin (median 32.459 peptides per experiment).

#### The PeptideAtlas tier system allows for easy analyses at different confidence levels

Proteins identified in PeptideAtlas builds are assigned to tiers based on the observed peptide evidence. Similar to a previous study^57^, we simplified it to four tiers (see Methods). 4,755 proteins were assigned to the highest tier (“canonical”), meaning that they have at least two uniquely mapping peptides. Of these, 1,246 are not part of the K-12 reference proteome, the most commonly used reference database for *E. coli* proteomics analyses. Peptides from 4,074 additional proteins were observed, but they could only be distinguished from other peptidoforms based on one uniquely mapping peptide (“marginally distinguished”).

The relative coverage varies across the selected reference proteomes (Fig 1C), with K-12 derivatives having the least percentage of unobserved proteins (around 5%), and pathogenic strains like NC101 and APEC the most (around 20%). As the K-12 MG1655 proteome was selected as the core proteome for the build, the number of canonical proteins for K-12 derivatives is higher due to proteins from this proteome given preference in case of ambiguity.

#### Strong experimental evidence for 1,376 proteins not observed until now

Of the 4,755 canonically identified protein isoforms in the *E. coli* PeptideAtlas build, 1,376 had no prior protein-level evidence in UniProt, even when accounting for evidence linked to homologous entries. Among these, 785 proteins originate from the well-studied K-12 MG1655 reference proteome. Notably, this set includes 358 proteins annotated as uncharacterized and an additional 541 proteins with limited existing knowledge, reflected by UniProt annotation scores below 3. For these poorly annotated MG1655 proteins, the PeptideAtlas resource offers valuable new evidence by indicating the specific strains and experimental conditions under which peptide support was observed, thereby providing a foundation for refining functional annotations and guiding future experimental investigations.

#### The webpage offers extensive information on PSM, peptide, protein, and proteome level

Upon accessing the build overview page, users are presented with summary statistics, including the total number of distinct peptides, post-translationally modified peptides, and protein identifications, alongside metrics such as the number of projects included and the search engine workflows applied. The interface allows intuitive navigation between the peptide, protein, and PSM levels, each of which provides layered insight into the *E. coli* proteome. On the protein level, users can explore proteins filtered by tiers, view supporting peptide evidence and from which experiments and strains they come from, determine the number and types of peptides matched, and assess evidence quality and coverage via visual tools such as protein coverage maps (Figure 2A). On the peptide level, the site offers overviews on peptide proteotypicity, its respective modification sites, and the number of PSMs supporting each peptidoform. Users can further interrogate PSMs, including raw spectrum annotations (Figure 2B) and quality metrics, aiding in the evaluation of peptide identification confidence. The interface also enables queries by protein accession and peptide sequence, and facilitates downloading of filtered or full identification lists for downstream analyses. Modifications can further be inspected interactively with PTMVision^64^, which has been extended to parse the Mass Modification Locations Table provided on the build page of each protein. PTMVision allows users to obtain an overview of PTMs on protein sequence and structure, and to inspect sites in more detail, e.g. with additional context from UniProt and residues in close contact to PTMs.

**Figure 2:**
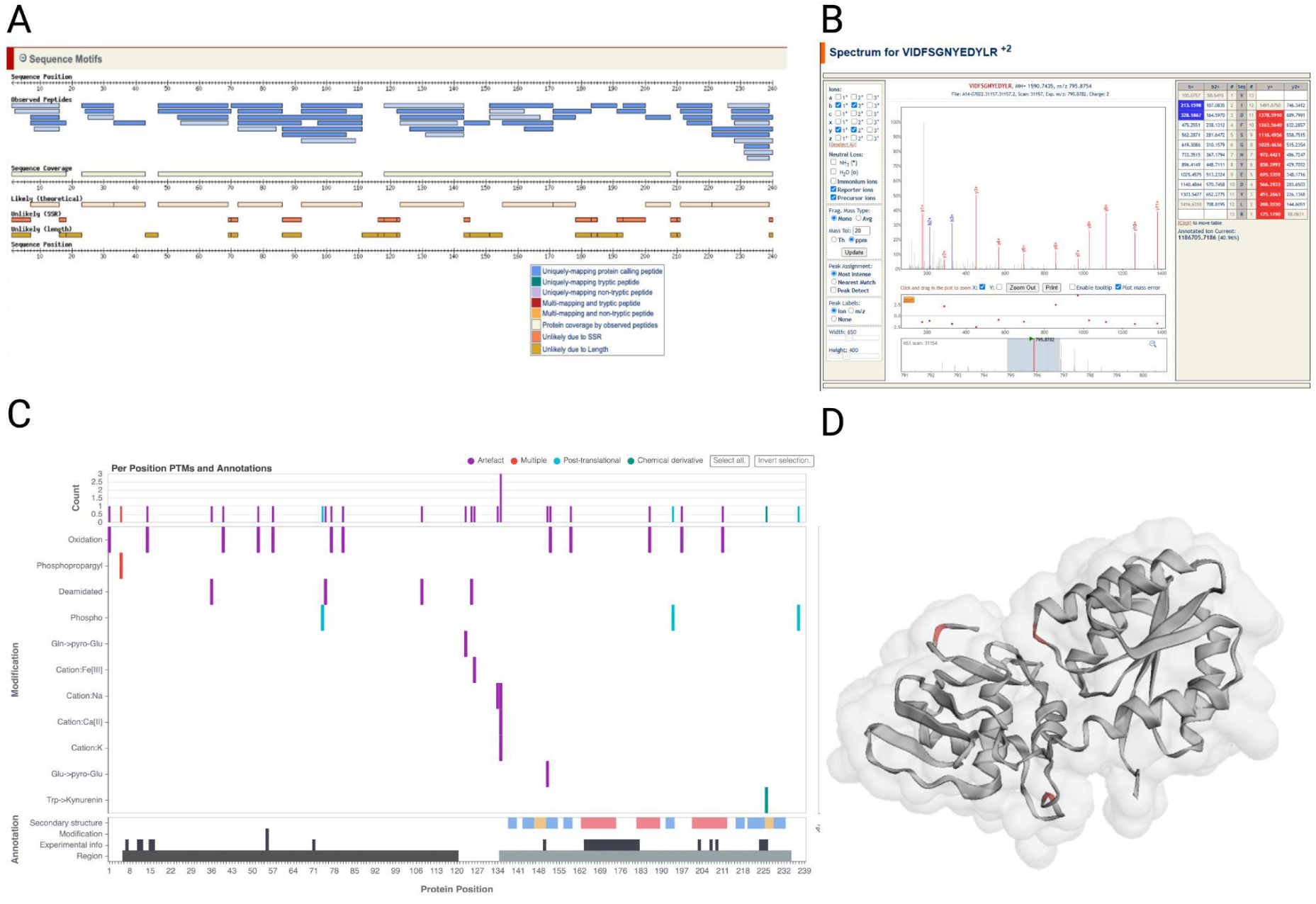
Exemplary functionalities of the E. coli PeptideAtlas build web interface. **A:** Alignment of observed peptides along the amino acid sequence for ompR (P0AA16), showing protein coverage with additional peptide information. **B:** Peptide evidence can be traced back down to the original spectrum, which can then be inspected in an annotated view. Example shown for one of eight spectra supporting the phosphorylation on S74 for ompR (USI mzspec:PXD008921:OR14_A3_20170411_MH_W3110_control_cipro_IMAC_2:scan:44656:SQS[Phospho]NPM PIIMVTAK/2). **C & D:** The protein modification site table can be uploaded to PTMVision to inspect modification sites interactively in both 2D (**C**) and 3D, with phosphorylations highlighted (**D**).

#### Bacteriophage peptides were detected in multiple projects

Despite strict efforts to maintain sterile conditions, bacteriophage contamination remains a major challenge in both industrial and laboratory settings, particularly during large-scale cultivation procedures^65^. In addition, certain *E. coli* strains naturally harbor prophage sequences within their genomes; among the strains examined, only DH5α was reported to contain five prophage sequences^66^. Peptides mapping uniquely to the phage proteins were detected in 17 out of 40 projects, with no observable preference for specific *E. coli* strains. Certain phages, such as T1, T2, T3, and T6, did not yield any detectable proteins at any confidence level. In contrast, T4 presented nine proteins with weak confidence and one with high confidence. For T7, three proteins were detected, including one canonical protein. Notably, 22 of 66 lambda phage proteins were identified with high confidence. An additional 11 proteins were identified with weak confidence, while two were marginally distinguished, and one protein remained indistinguishable from other phage proteins. Furthermore, peptide evidence for 16 of the 22 canonical lambda phage proteins was found in one specific dataset (PXD041301, strain DSM 105380) that included samples enriched for outer membrane vesicles. This preparation method likely enhances the detection of phage proteins because outer membrane vesicles are a defense mechanism of gram negative bacteria and work as “decoys” that carry membrane receptors that bacteriophages bind to^67^.

### The predicted and the observed *E. coli* pan-proteome

To gain a comprehensive view of the *E. coli* pan-proteome, we clustered proteins from selected reference proteomes using varying similarity thresholds, allowing us to classify protein families into core, accessory, and strain-specific orphan groups. This approach provides insight into the structural organization of the pan-proteome and its relevance across diverse *E. coli* strains. By linking these clusters to peptide identifications from the PeptideAtlas build, we further evaluated which parts of the proteome are readily detectable by mass spectrometry and which remain undetected. Our analysis demonstrates that many proteins go unobserved due to physicochemical properties that hinder their identification, such as small size, limited tryptic peptide yield, and membrane localization. Nonetheless, the use of diverse enrichment and fractionation strategies enabled the detection of a broad range of proteins, including those typically difficult to identify, and revealed strain-specific candidates with potential utility for typing and functional characterization.

#### Homology clustering and pan-proteome composition

Clustering the proteins included in the selected reference proteomes at varying sequence similarity thresholds revealed a clear, and expected trend in the relationship between the number of homology clusters and their mean size (Figure 3A). As the similarity threshold decreased, the number of clusters diminished, while the mean cluster size increased, indicating greater aggregation of homologous sequences. At a 70% sequence similarity threshold, the pan-proteome, consisting of 27,806 distinct proteoforms, was divided into 8,310 homology clusters, with an average of 3.3 proteoforms per cluster. This suggests that, on average, each *E. coli* protein has at least three distinct proteoforms; the distributions are shown in Supplementary Figure 2. The homology clusters were further classified into core, accessory, and orphan categories (Figure 3B). Of these, 2,871 clusters (35%) were present in only one proteome, representing orphan proteins, while 2,649 clusters (32%) appeared in two or more proteomes but not in all, indicating accessory proteins. The remaining 2,790 clusters (33%) were present across all proteomes, representing core proteins.

**Figure 3:**
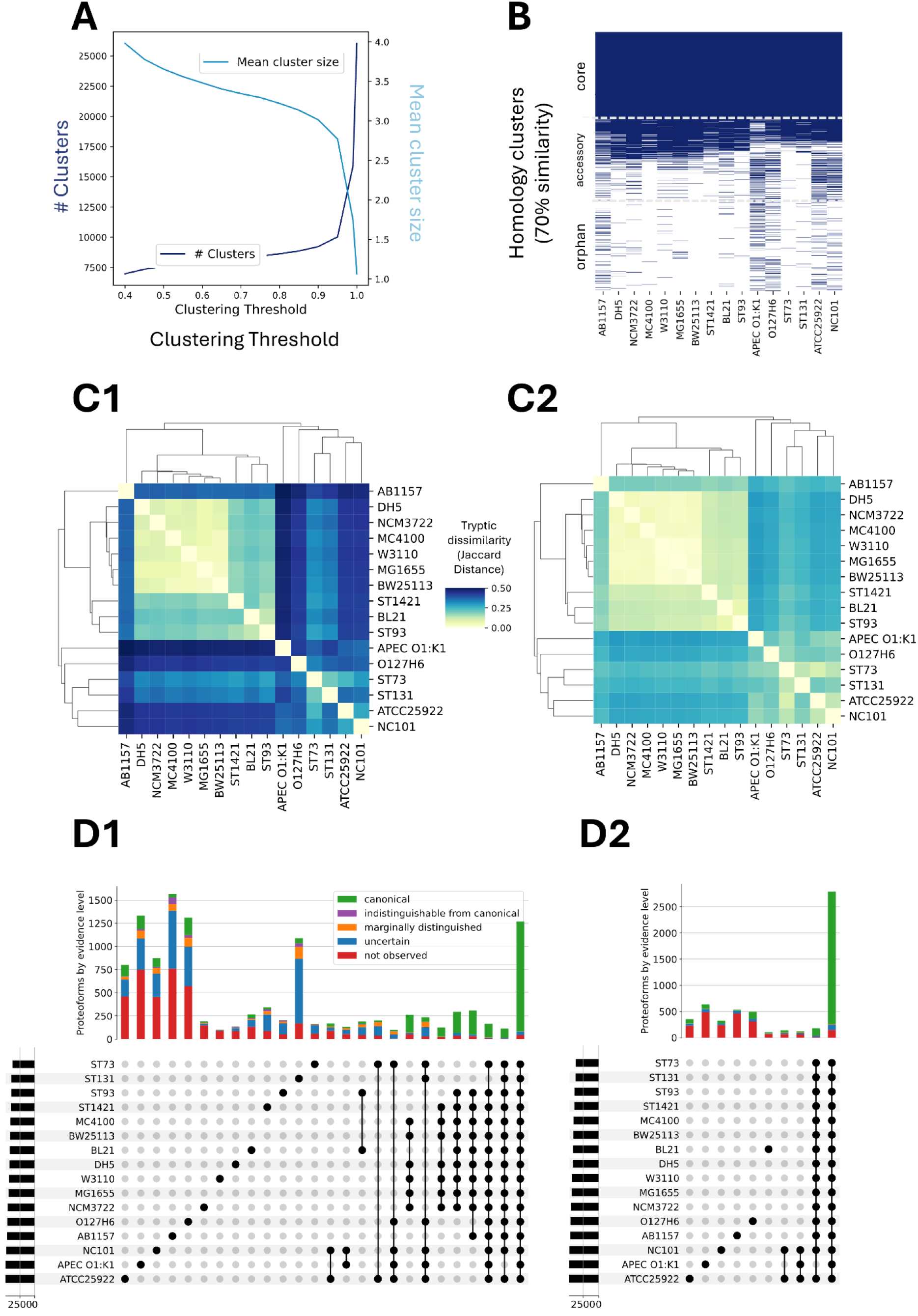
Predicted and observed E. coli Pan-proteome. **A:** Number of homology clusters and their sizes as a function of sequence similarity threshold used for clustering. **B:** Homology clusters of E. coli Pan-proteome after clustering at 70% sequence similarity. **C:** Pairwise Jaccard distances of tryptic digests of UniProt proteomes (C1) vs. pairwise Jaccard distances of tryptic digests of UniProt proteomes only considering peptides observed in the PeptideAtlas build (C2). **D:** Upset plot of proteome clustering at 99% (D1) and 70% (D2) sequence similarity. Colors indicate highest level of evidence of all proteins in the cluster. Only sets with more than 100 protein clusters are shown.

At a 99% similarity threshold, the core proteome is significantly smaller than at 70%, with only 1,269 proteins common across all strains (Supplementary Figure 3). Only 247 proteins are shared with identical sequences across all strains.

#### Peptide similarity and observability in proteomics datasets

The pairwise Jaccard distances of tryptic digests of UniProt proteomes (Figure 3C1) revealed distinct clustering patterns among *E. coli* strains. For example, the K-12 derivatives (MC4100, BW25113, DH5, W3110, MG1655, and NCM3722) are clustered together, while the proteomes from the pathogenic *E. coli* (e.g. NC101, APEC) show larger differences to all other strains.

When considering only the peptides observed in the PeptideAtlas build (Figure 3C2), the similarity between strains increased. This observation may be attributed to two factors. First, the limitations of experimental peptide coverage, as proteomic experiments typically detect only a subset of proteins, despite computationally predicted proteomes offering a more comprehensive overview. Second, sequence variations present in the proteomes may not have been represented in the analyzed samples, or may have been overlooked by the search engines during peptide identification.

#### Comparison of proteome clustering at high and low stringency

The proteome clustering at high (99%) and moderate (70%) sequence similarity thresholds exhibited distinct patterns (Figure 3D). The UpSet plot for 99% similarity (Figure 3D1) showed a greater number of unique protein clusters, indicative of strain-specific proteoforms. At 70% similarity (Figure 3D2), more clusters merged, emphasizing broader functional conservation. The color coding of protein clusters based on their level of evidence demonstrated that a significant fraction of the proteome has strong experimental support, while some clusters remained uncertain or unobserved.

The proteomes of strains ATCC 25922, APEC, NC101, AB1157, O127H6, and ST131 contain the highest numbers of unique proteoforms or proteins at 99% similarity (Figure 3D1). These numbers decrease significantly when the similarity threshold is slightly lowered, indicating that many of these proteoforms are from proteins shared across strains. However, some strain-specific clusters are retained even at lower thresholds (around 300-600 proteins, Figure 3D2), suggesting the presence of unique proteins for certain strains. While most of these strain-specific proteins lacked sufficient peptide evidence, a small subset was more confidently observed. These latter strain-specific proteins could potentially serve as markers for strain typing. For example, in the case of NC101, a pathogenic strain associated with adherent-invasive *E. coli* (AIEC) that currently lacks molecular markers^68^, 56 strain-specific proteins were identified with canonical evidence, 23 of which were exclusively observed in NC101 strains. One such protein, cdiI, is part of the Contact-Dependent Inhibition (CDI) system, which is present in several *E. coli* strains, but the version in NC101 (Identifier UPI0001DBCD95 / CDII_ECONC) differs significantly from the others. This protein was identified with 69.8% coverage and five unique peptides, all exclusively from NC101 samples. Given its strain specificity, its high detectability by mass spectrometry, and its lack of peptide matches in other strains, cdiI represents a promising candidate for strain typing.

The proteomes of pathogenic strains and isolates (ST73, NC101, APEC, ATCC25922) also show an overlap in proteins at 99% similarity. Their overlap could indicate conserved proteins necessary for virulence or survival in host environments. For the K-12 derivatives (MC4100, BW25113, DH5, W3110, MG1655, NCM3722), proteomes have a large overlap due to close genetic relationships.

Independent of the chosen similarity threshold, confident peptide evidence was observed for at least one protein in most of the core homology clusters. However, for a small fraction, no evidence was found. As these proteins are highly conserved across strains, they are probably important for basic functionalities and therefore present in the analysed samples, but missed due to technical limitations. In the following, we explore these limitations.

#### The dark part of the pan-proteome is mostly dark due to technical limitations

For 6,445 out of 27,806 (23%) proteoforms (or for 3,276/8,310 protein clusters at 70% similarity), no peptide evidence was observed in the *E. coli* PeptideAtlas. To investigate the effects of technical limitations and biases in the build, we characterized the observed and unobserved proteins by multiple characteristics: protein mass, length, number of tryptic peptides, cellular localisation, and isoelectric point. To facilitate this analysis, we defined the observed pan-proteome as the set of proteoforms that have peptide evidence in the PeptideAtlas, independent of their tier.

Analysis of key properties revealed that a significant portion of the pan-proteome’s proteoforms is likely unobserved due to their chemical properties, which are not ideal for mass spectrometry analysis. Unobserved proteoforms predominantly have higher isoelectric points (Figure 4A), are mostly smaller, and give rise to less tryptic peptides that can be identified in mass spectrometry experiments (Figure 4B). Many of the unobserved proteoforms are also predicted to reside in the cytoplasmic membrane (Figure 4C), suggesting that they are primarily hydrophobic. This presents a challenge for detection, as hydrophobic proteins are typically less compatible with sample processing, protein isolation, and mass spectrometry due to their low solubility and low tryptic peptide yield.

**Figure 4:**
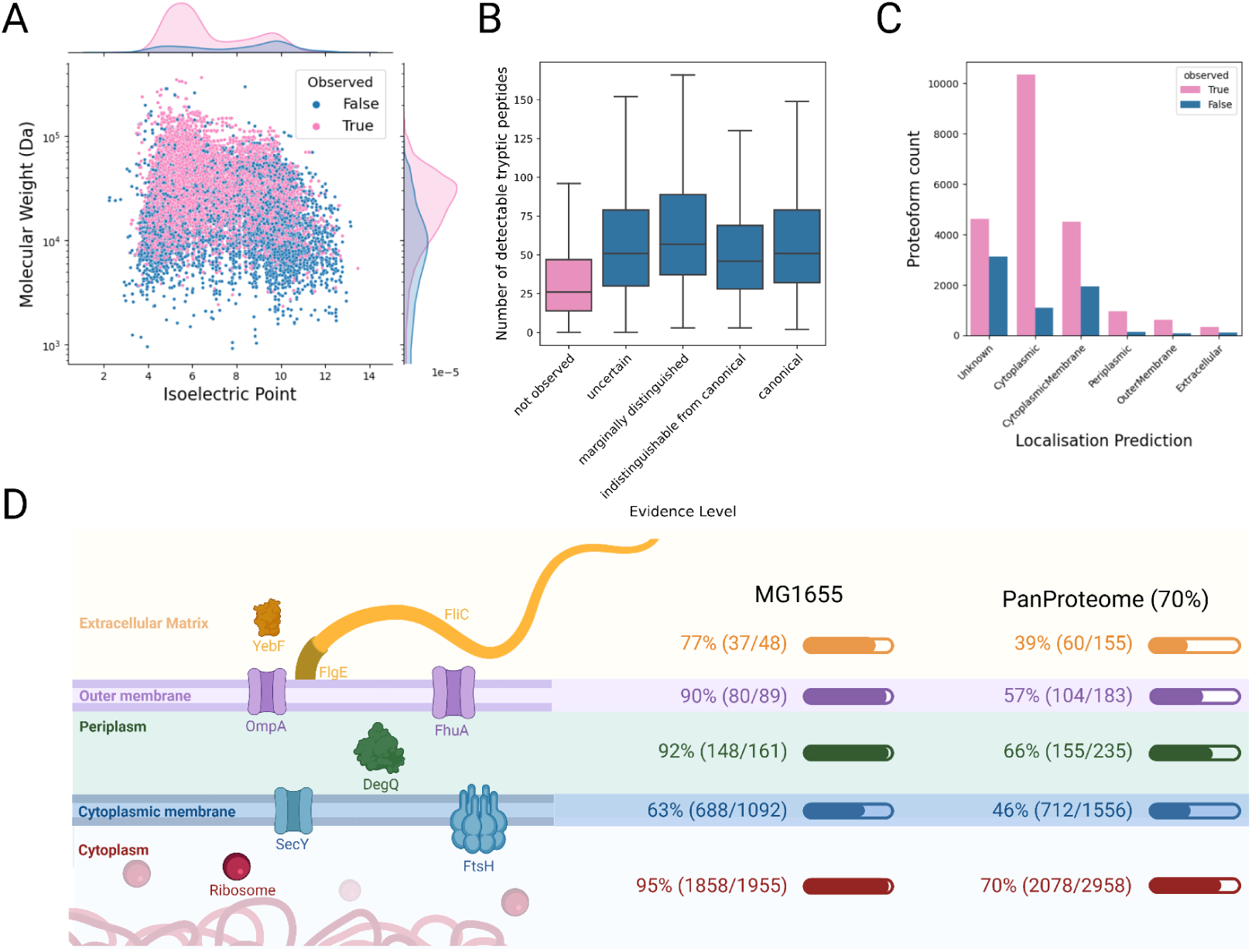
Properties of the observed vs. the unobserved Pan-proteome. **A:** Distribution of proteoform weights and isoelectric points of observed vs. unobserved proteoforms. **B:** Distributions of number of detectable tryptic peptides across evidence tiers. **C:** Number of observed vs. not observed proteoforms across cellular localisations as predicted by PSORTb. **D:** Cellular locations of the proteoforms in E. coli cells. Left: Structure of the E. coli cell membrane system with example proteins. Right: Identification proportion with canonical evidence in the E. coli K-12 MG1655 proteome, and the pan-proteome clustered at 70% similarity. For 3,224 homology clusters, and for 1,059 MG1655 proteins, no localisation could be predicted by PSORTb.

Despite these challenges, we were able to identify membrane proteins both in the cytoplasmic membrane and in the outer membrane (Figure 4D). This is likely thanks to the combination of fractionation and enrichment techniques and different sample preparation methods. For example, probe sonication has been shown to be effective for processing membrane proteins in bacterial systems^41^. In addition, even proteins that are generally less amenable to mass spectrometry, such as highly hydrophobic or membrane-associated proteins, may still contain accessible domains or loop regions that yield one or more detectable peptides, facilitating their partial observation under optimal conditions.

### Post-translational modifications in *E. coli*

PTMs are essential regulatory mechanisms in all organisms, including bacteria, and they play a crucial role in the control of protein function, stability, localization, and interactions. In bacteria, PTMs can modulate a variety of processes such as metabolism, stress response, antibiotic resistance, and virulence. As highlighted by Macek *et al*.^69^, PTMs in bacteria have gained significant attention due to their involvement in essential cellular processes, such as signal transduction, energy production, and response to environmental changes.

In our reprocessing of large-scale *E. coli* MS data, a wide variety of PTMs were detected. A total of 113,258 modification events were identified across the datasets, corresponding to multiple PTM types. These modification events (composed of protein residue and the modification type, e.g. Tyr23-phosphorylated) were categorized by their biological relevance, as per the Unimod classification^70^. The most observed classes are *artefacts* (41,353 events), *isotopic labels* (32,005 events), *chemical derivatives* (21,628 events), and *post-translational modifications* (10,887 events) (Figure 5A). 830 modification events were assigned to more than one Unimod class, these are grouped under “Other”.

**Figure 5:**
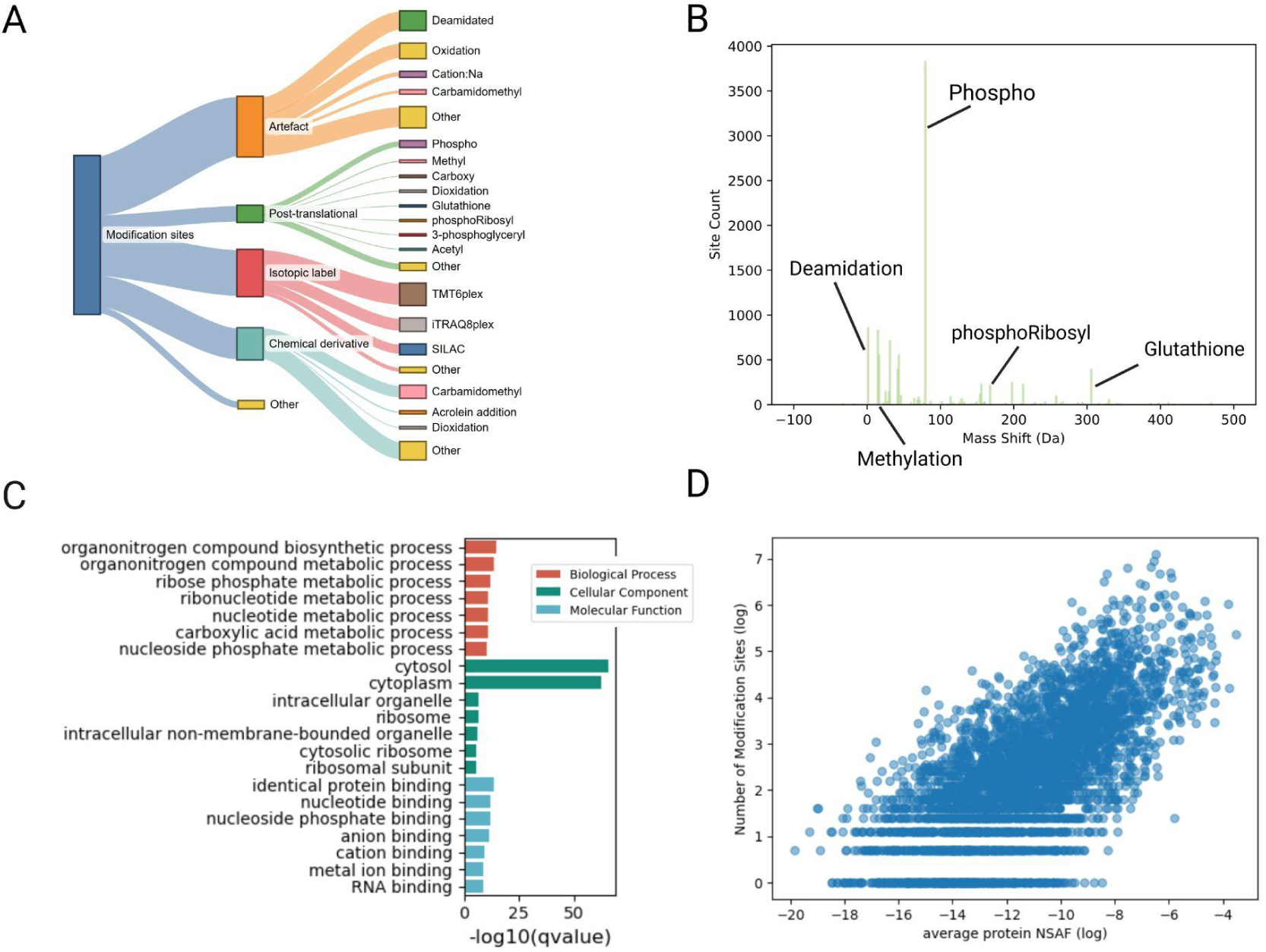
PTMs identified in the E. coli Pan-proteome build. **A:** Identified modifications and their classification according to Unimod. **B:** Distribution of mass shifts induced by identified post-translational modifications, showing number of modification sites on y-axis. **C:** Significantly enriched GO Terms of glutathionylated proteins. Colors indicate different GO Term classes. **D:** Plotting protein abundance (NSAF) against number of modification sites shows that peptides from higher abundant proteins are more often identified in modified forms.

Artefacts comprise modifications that arise as byproducts of sample preparation, such as methionine oxidation resulting from exposure to atmospheric oxygen or reactive oxygen species during digestion and handling. The most prevalent artefactual modifications are deamidation (12,626 sites) and oxidation (9,899 sites), primarily affecting asparagine and glutamine residues, and methionine residues, respectively. In contrast, chemical derivatives represent modifications introduced intentionally for experimental purposes, with carbamidomethylation being the most common (8,115 sites), typically targeting cysteine residues during alkylation.

The selected projects for the build include four TMT, three SILAC, and one iTRAQ experiment, which is reflected in the number of identified isotopic labels. Other groups include glycosylations, nonclassified modifications, and co- or pre-translational modifications. From a biological perspective, modification events classified as post-translational modifications are the most interesting. Here, the most prominent modification type is phosphorylation (3,806 sites) due to the integration of phosphoenriched samples, followed by deamidation (854 sites), methylation (719 sites), and carboxylation (554 sites).

Among the identified PTMs, phosphorylation was found on the highest number of residues, with 3,806 detected modification sites. Phosphorylation is one of the most studied and important PTMs in both eukaryotes and prokaryotes. In bacteria, phosphorylation plays a critical role in regulating signal transduction pathways, cell cycle progression, metabolism, and response to environmental changes. In *E. coli*, many key proteins involved in stress responses, signal transduction (e.g., two-component systems), and metabolic regulation are known to be phosphorylated. Phosphorylation often occurs on serine, threonine, and tyrosine residues. Our data indeed reveals a substantial presence of phosphorylation at serine (1,753 sites), threonine (996 sites), and tyrosine (553 sites). It has been previously described in the literature that phosphorylation sites are enriched in disordered regions of proteins in humans^71,72^. In *E. coli,* we also observed more phosphorylated sites in disordered regions as compared to nonphosphorylated residues, however the difference was rather small (Supplementary Figure 5).

Glutathione (GSH) is a critical antioxidant in bacterial cells, maintaining the redox balance by scavenging reactive oxygen species (ROS) and other free radicals. The modification of proteins by glutathione, referred to as glutathionylation, has been implicated in redox regulation, cellular protection against oxidative stress, and the regulation of key enzymes in *E. coli*. We identified 253 glutathione sites on cysteine residues. The identification evidence mainly derives from a project in which the bacteria were subjected to ciprofloxacin-induced antibiotic stress (PXD050358: 4582/6185 glutathionylated PSMs). Metal ion binding proteins were significantly (adjusted p-value < 0.05) overrepresented in glutathionylated proteins (Figure 5C), underlining the role of glutathionylation in cellular defense mechanisms, particularly in the homeostasis and resistance to metal ions^73^.

Phosphoribosylation, the modification of proteins by the addition of a phosphoribosyl group, is another important PTM in bacteria, particularly in the context of nucleotide metabolism and secondary metabolism. In our data, phosphoribosylation was found on glutamic acid (125 sites) and aspartic acid (94 sites), with some occurrences on arginine (7 sites). This modification is most likely involved in bacterial processes related to nucleotide biosynthesis and regulation, as phosphoribosylation is a critical step in purine and pyrimidine biosynthesis pathways. The presence of phosphoribosylation in *E. coli* may be linked to the regulation of metabolic pathways that are essential for cellular growth and proliferation, especially under varying environmental conditions.

We further investigated the potential bias of PTM site identifications. As modifications are thought to exist mostly at substoichiometric levels^74^, it is likely that we will identify PTMs on proteins that are higher abundant to begin with. Indeed, the number of modification sites increases with protein abundance (Figure 5D).

We investigated whether the observed patterns of post-translational modifications could be explained by the available experimental metadata. Principal clustering of the runs was primarily driven by project affiliation, indicating pronounced batch effects as a major contributor to variation in PTM profiles (Supplementary Figure 6A). Clustering by strain was also observed (Supplementary Figure 6B); however, due to the strong correlation between strain and experimental batch, it remains challenging to disentangle the respective contributions of biological versus technical factors. No clear clustering was detected based on other metadata variables such as growth medium composition (Supplementary Figure 6C and 6D), optical density, or other culture conditions. While this absence of signal does not exclude potential biological influence on PTMs, it is likely that such effects are obscured by dominant batch effects or limited by insufficient annotation coverage for certain variables.

## Discussion & conclusion

In this study, we have presented a comprehensive reprocessing and analysis of large-scale *Escherichia coli* proteomics data from multiple global research projects. By employing both closed (TPP with MSFragger) and open (ionbot) search strategies, we have integrated data from 40 selected projects, resulting in the identification of 4,755 high-confidence proteins. These identifications represent a significant contribution to the understanding of *E. coli*’s proteome under a variety of conditions, including stress responses, antibiotic exposure, and genetic modifications. By using a core proteome derived from the well-annotated K-12 MG1655 strain and combining it with additional strains and phages, we have created a detailed *E. coli* PeptideAtlas, which serves as a valuable resource for future studies. It allows for the visualization and exploration of protein identifications across strains, with detailed annotations on coverage, abundance, and PTMs. Additionally, we detected evidence for previously unidentified proteins, providing strong experimental evidence for 1,376 canonical proteins without protein evidence in UniProt.

To ensure compatibility with both search strategies, our analysis was restricted to datasets acquired using DDA, with the exception of one DIA dataset, which was processed using only the TPP. This choice was motivated by the current limitations in open search capabilities for DIA data, as ionbot does not support this acquisition mode. Although DIA holds promise for improved reproducibility and depth of coverage, the inability to perform open modification searches on DIA data remains a constraint. We anticipate that as search engines evolve to accommodate open searches in DIA mode, future versions of this PeptideAtlas resource may be expanded to include such datasets, thereby further enhancing proteome coverage and biological insight.

As we identified phage proteins in multiple runs, we suggest including bacteriophage proteins in the search space of database searches to improve completeness. Despite stringent efforts to maintain sterile conditions, bacteriophage contamination remains a major challenge. The inclusion of phage sequences in the database facilitates the detection of phage-related peptides, improving the spectrum identification rate and making it easier for researchers to account for and identify potential phage contamination and prophages in their studies. This approach is particularly valuable in settings where the likelihood of phage protein presence is elevated, such as laboratories that routinely handle bacteriophages, studies involving newly uncharacterised strains, or experiments that enrich for outer membrane vesicles. Moreover, incorporating these sequences can aid in improving false discovery rate (FDR) control by reducing the number of incorrectly assigned matches to bacterial proteins when spectra in fact originate from phage-derived peptides.

While there is significant progress in making data publicly available, as evidenced by the growing number of public datasets on platforms like ProteomeXchange, a key challenge remains in the provision of metadata annotation. Missing or inconsistent metadata can hinder the reuse of datasets, preventing these from reaching their full potential. Improved and standardized metadata annotation would greatly enhance the ability of researchers to accurately interpret the data and facilitate its integration into larger-scale studies, promoting better reproducibility and broader utilization of publicly available data.

To conclude, this work provides a high-quality, deeply annotated, and easily accessible resource that significantly enhances our understanding of the *E. coli* proteome across a wide range of biological contexts. By systematically reprocessing diverse datasets and integrating robust search strategies, we have built a PeptideAtlas that not only expands the coverage of known proteins, including previously unconfirmed ones, but also enables exploration of proteomic variation across strains and conditions. The PeptideAtlas build (called 2024-11 ionbot) is available online and includes all identified peptide-spectrum matches, post-translational modifications, and associated metadata. This comprehensive resource supports targeted mass spectrometry assay development, PTM-focused studies, and facilitates comparative analyses across strains. Ultimately, it represents a valuable foundation for future *E. coli* proteomics research, fostering data reuse, reproducibility, and discovery.

## Data availability

The PeptideAtlas build is available online at https://db.systemsbiology.net/sbeams/cgi/PeptideAtlas/buildDetails?atlas_build_id=585. Code for producing the figures and the search fastas are available on GitHub (https://github.com/CompOmics/ecoli-peptideatlas-manuscript) and Zenodo (https://doi.org/10.5281/zenodo.15551248). The mass spectrometry proteomics data and associated metadata have been deposited to the ProteomeXchange Consortium via the PRIDE^75^ partner repository with the dataset identifier PXD058808. Modification Sites are provided in Supplementary Table 3.

## Competing Interests

The authors declare no competing financial interests.

## Author Contribution statement

C.J.: Investigation, Data Curation, Writing - Original

Draft. A.H.: Investigation.

F.A.: Investigation.

C.C.: Resources and Project administration - Review & Editing. E.D.: Methodology.

J.Z.: Formal analysis. Z.S.: Formal analysis. K.V.: Formal analysis.

R.B.: Writing - Review & Editing.

K.N.: Supervision, Funding acquisition - Review & Editing.

L.M.: Supervision, Funding acquisition - Review & Editing.

R.G.: Supervision, Writing - Review & Editing.

T.V.: Supervision, Writing - Review & Editing.

## Acknowledgements

C.J. has received funding from the European Union’s Horizon Europe under grant agreement no. 101119980. L.M. acknowledges funding from the Horizon Europe Projects BAXERNA 2.0 [101080544] and COMBINE [101191739], and from the Ghent University Concerted Research Action [BOF21/GOA/033]. L.M. and C.C. are further supported by the CHIST-ERA project ODEEP-EU [G0GDV23N] and FWO SRN [W005325N]. R.B., L.M. and R.G. acknowledge funding from the Research Foundation Flanders (FWO) [12A6L24N, 1SE3724N, G010023N, G028821N, 12B7123N]. C.C. and L.M. acknowledge CNRS and the Agence Nationale de la Recherche via the French Proteomic Infrastructure (ProFI UAR2048; ANR-10-INBS-08-03 and ANR-24-INBS-0015).

## Supplementary Figures

**Supplementary Figure 1:**
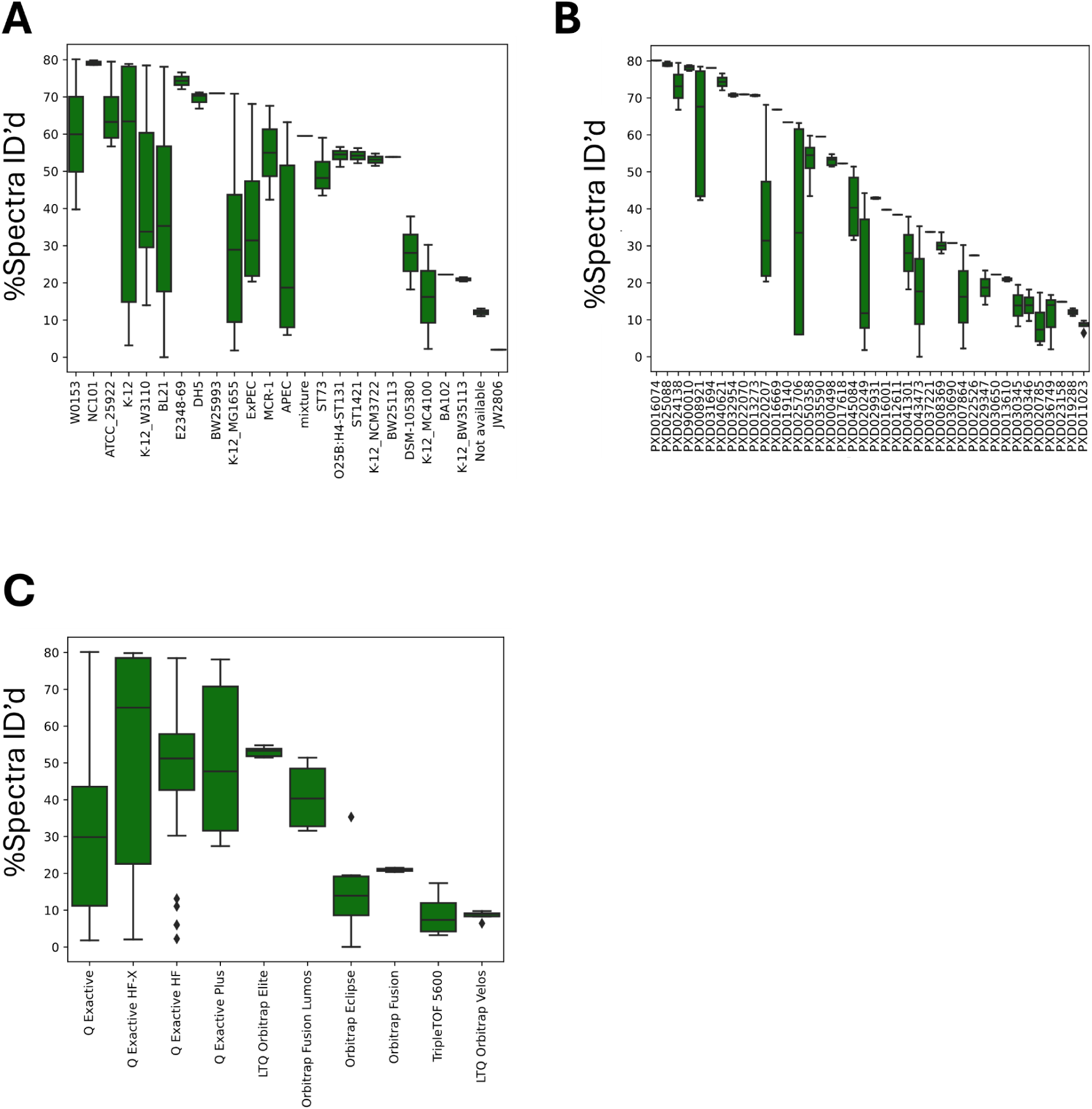
Differences in experiment level spectrum matching rates between strains (**A**), projects (**B**), and mass spectrometer instruments (**C**).

**Supplementary Figure 2:**
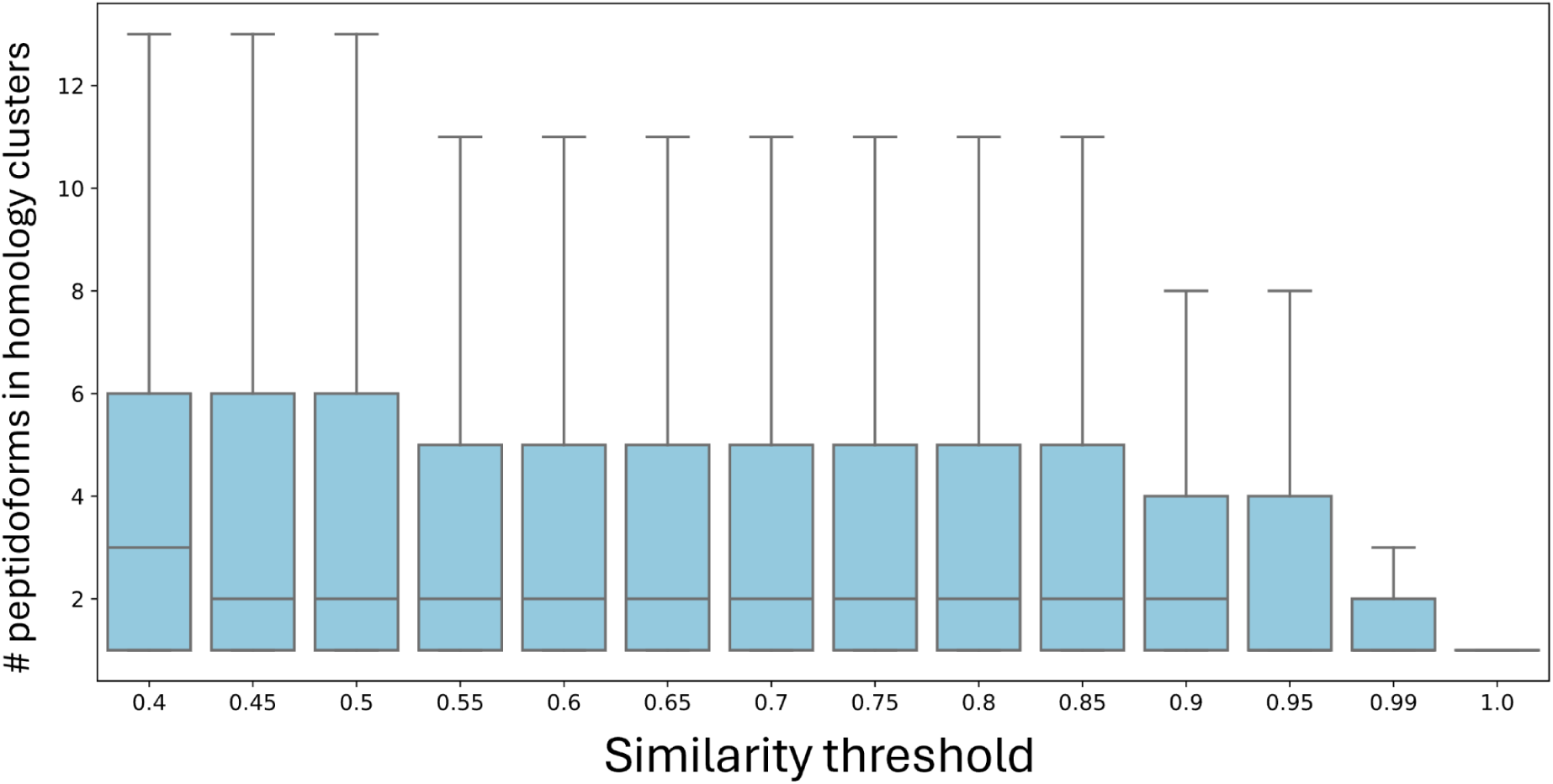
Distribution of number of peptidoforms per homology cluster, as a function of similarity threshold.

**Supplementary Figure 3:**
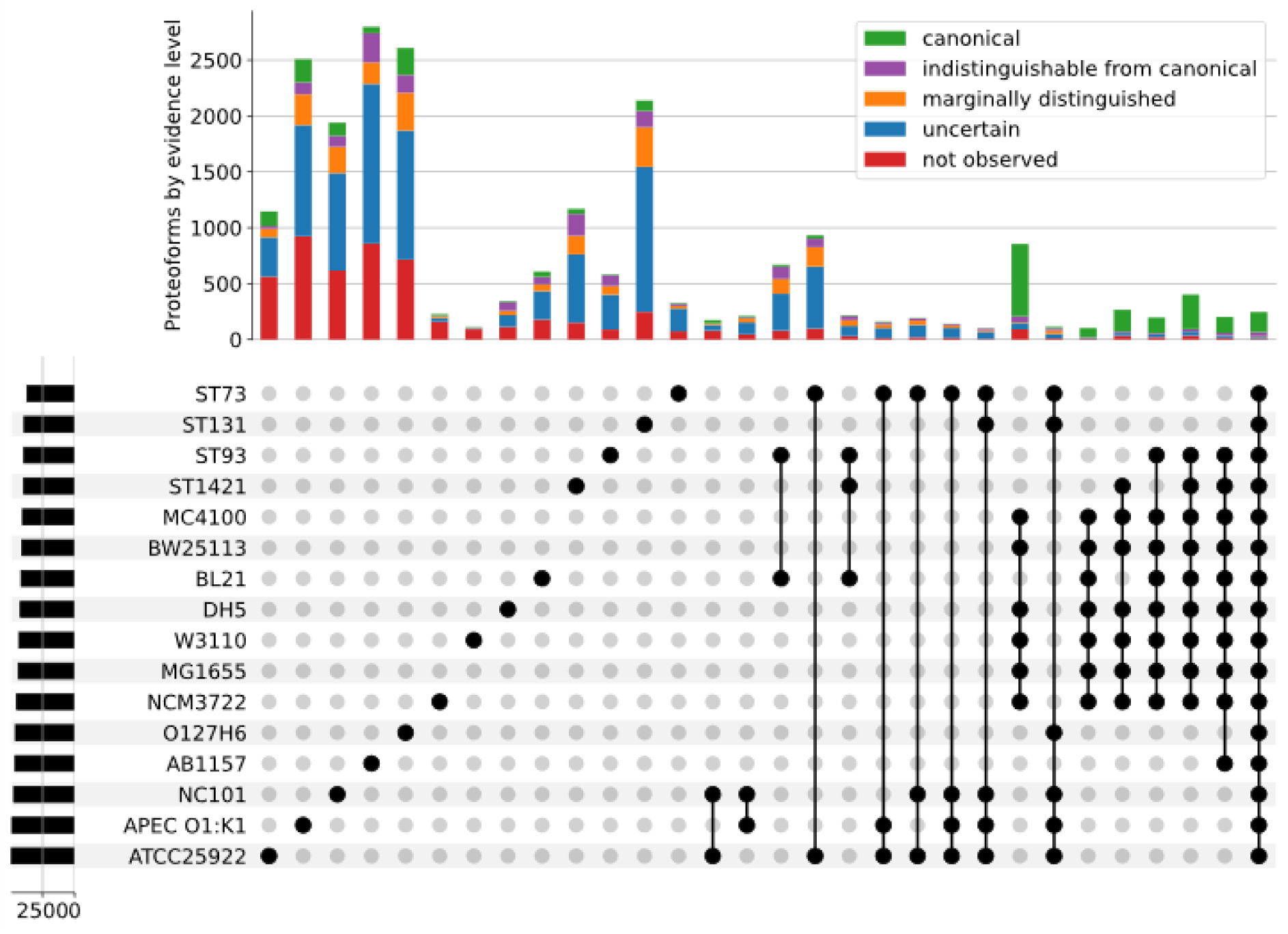
Pan-proteome clustering with CD-HIT at 100% similarity.

**Supplementary Figure 4:**
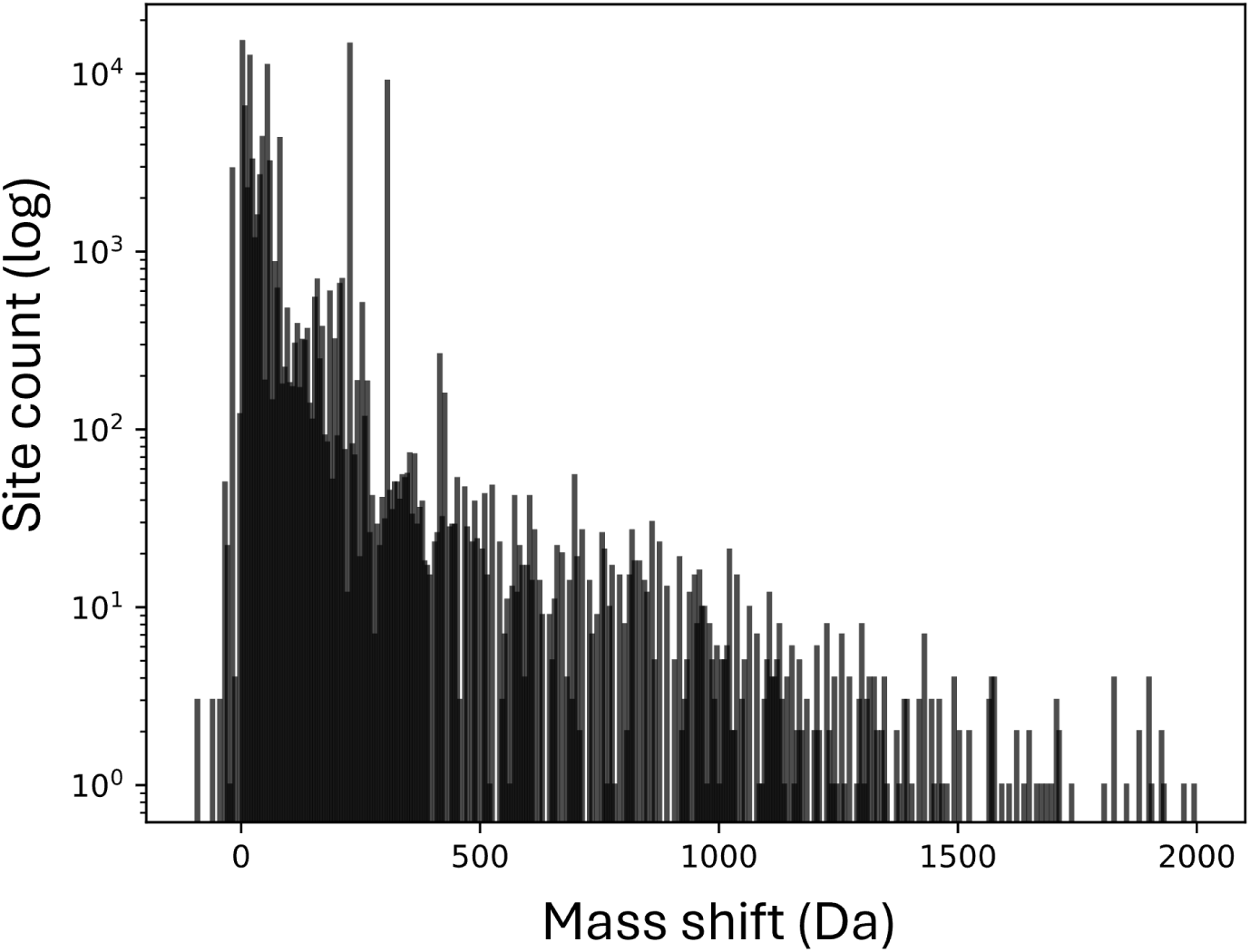
Histogram of mass shifts induced by modifications (including all Unimod classes) identified in the build uniquely mapping to canonical proteins.

**Supplementary Figure 5:**
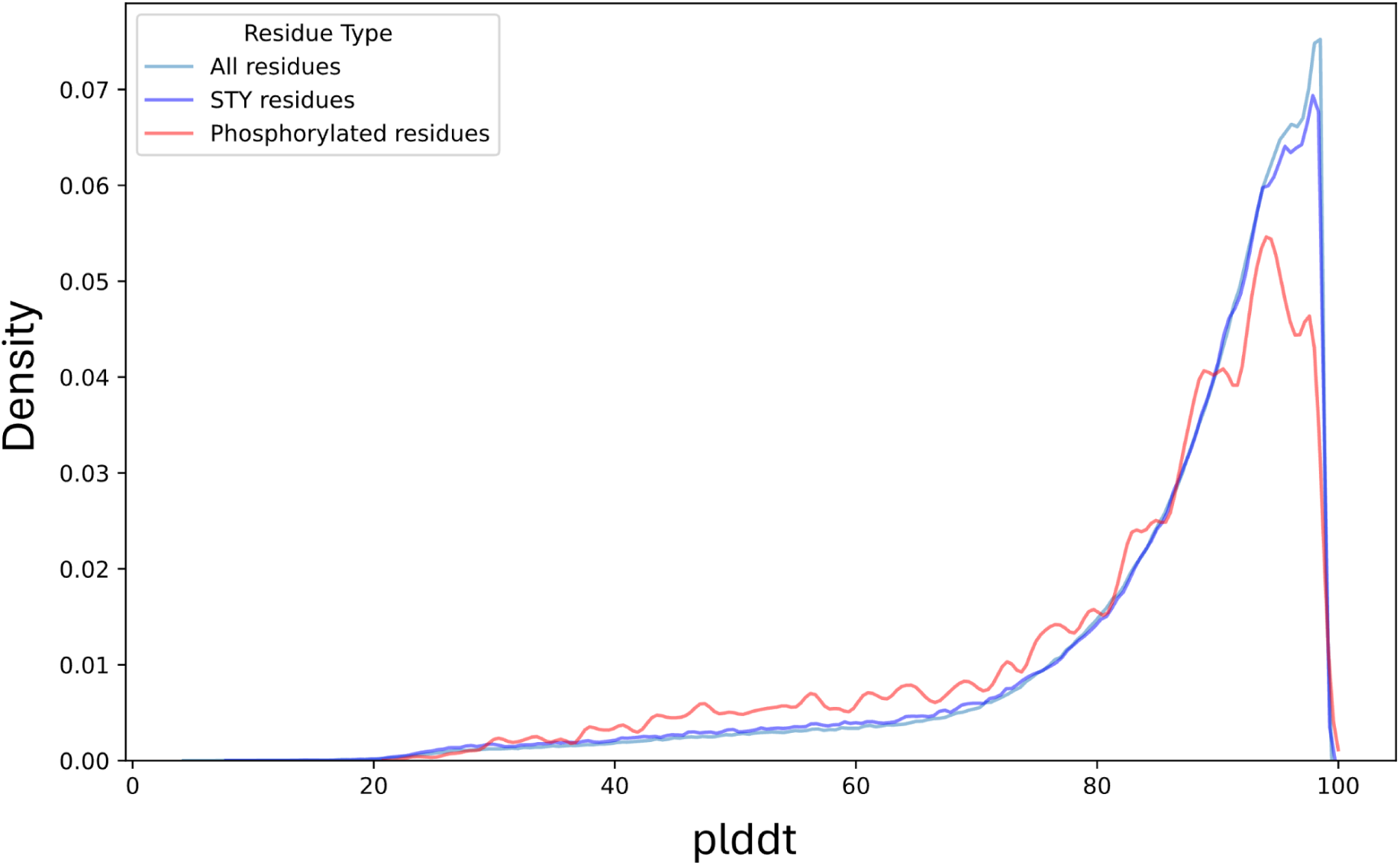
Distribution of ESMFold pLDDT values of all residues (bright blue), STY residues (dark blue), and phosphorylated residues (red). Data is based on ESMFold predictions of proteins in the highest confidence tier, and only phosphorylation sites matching uniquely to a canonical protein were included.

**Supplementary Figure 6:**
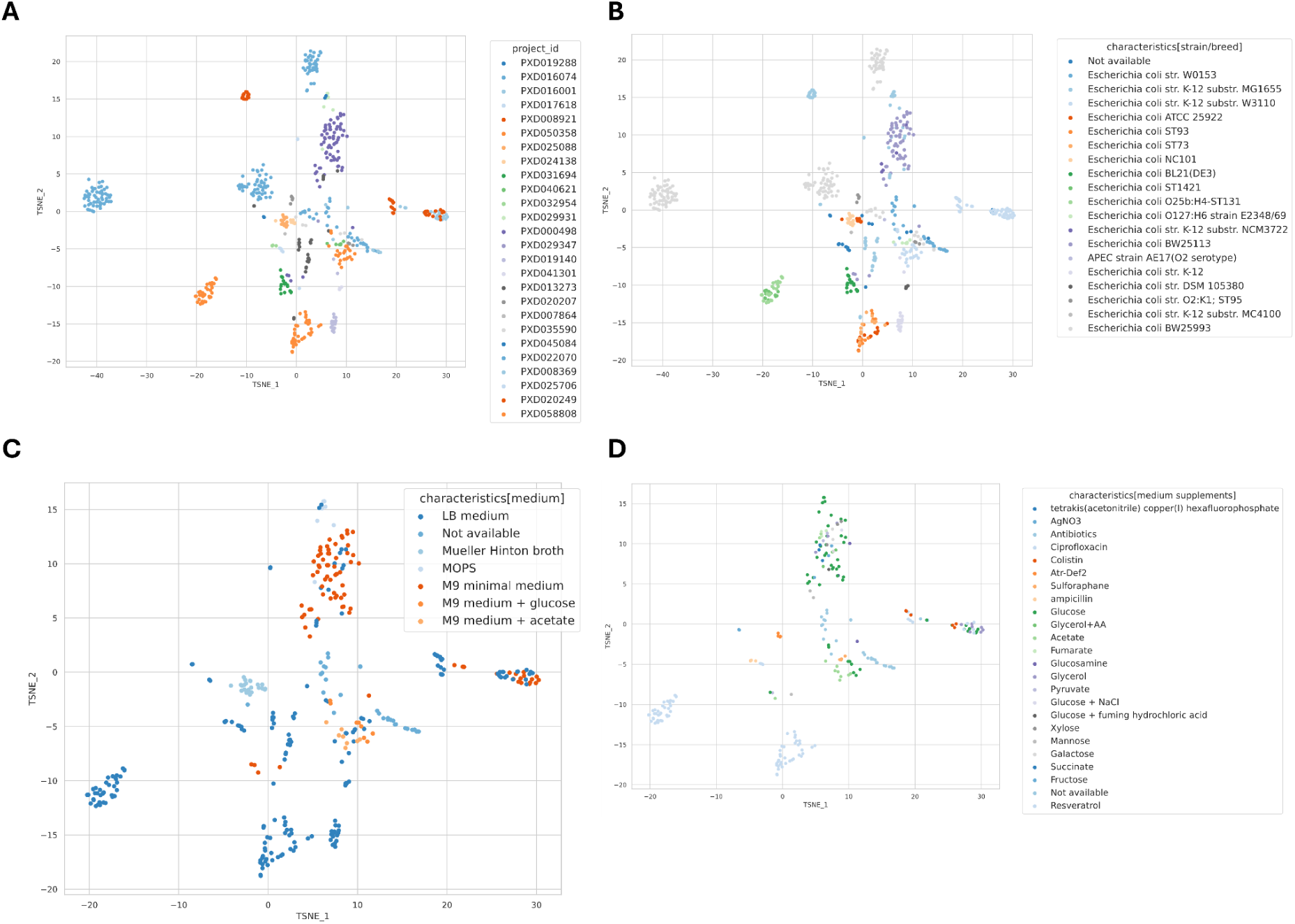
t-SNE of all runs according to post-translational modifications, colored by experiment (**A**), medium (**B**), strain (**C**), and medium supplements (**D**). t-SNE based on the observation counts for each PTM per run, i.e. each run is represented by a vector specifying how many times each PTM was found in that run. Note that runs without any observed modifications classified as post-translational are not represented.

## Notes

https://zenodo.org/records/15551249

### Competing Interest Statement

The authors have declared no competing interest.

## References

1. Geurtsen, J. et al. Genomics and pathotypes of the many faces of *Escherichia coli*. FEMS Microbiol. Rev. 46, fuac031 (2022).

2. Kaper, J. B., Nataro, J. P. & Mobley, H. L. T. Pathogenic Escherichia coli. Nat. Rev. Microbiol. 2, 123–140 (2004).

3. Touchon, M. et al. Organised Genome Dynamics in the Escherichia coli Species Results in Highly Diverse Adaptive Paths. PLoS Genet. 5, e1000344 (2009).

4. Midha, M. K. et al. A comprehensive spectral assay library to quantify the Escherichia coli proteome by DIA/SWATH-MS. Sci. Data 7, 389 (2020).

5. Schmidt, A. et al. The quantitative and condition-dependent Escherichia coli proteome. Nat. Biotechnol. 34, 104–110 (2016).

6. Yu, Y. et al. Predictive Signatures of 19 Antibiotic-Induced *Escherichia coli* Proteomes. ACS Infect. Dis. 6, 2120–2129 (2020).

7. Rudenko, I., Ni, B., Glatter, T. & Sourjik, V. Inefficient Secretion of Anti-sigma Factor FlgM Inhibits Bacterial Motility at High Temperature. iScience 16, 145–154 (2019).

8. Liu, Y. et al. Effect of spaceflight on the phenotype and proteome of Escherichia coli. Open Life Sci. 18, 20220576 (2023).

9. Deutsch, E. W. et al. Trans-Proteomic Pipeline: Robust Mass Spectrometry-Based Proteomics Data Analysis Suite. J. Proteome Res. 22, 615–624 (2023).

10. Degroeve, S. et al. Ionbot: A Novel, Innovative and Sensitive Machine Learning Approach to LC-MS/MS Peptide Identification. http://biorxiv.org/lookup/doi/10.1101/2021.07.02.450686 (2021) doi:10.1101/2021.07.02.450686.

11. Jensen, K. F. The Escherichia coli K-12 ‘wild types’ W3110 and MG1655 have an rph frameshift mutation that leads to pyrimidine starvation due to low pyrE expression levels. J. Bacteriol. 175, 3401–3407 (1993).

12. Schmidt, A. et al. The quantitative and condition-dependent Escherichia coli proteome. Nat. Biotechnol. 34, 104–110 (2016).

13. Rowland, E. A. et al. Sirtuin Lipoamidase Activity Is Conserved in Bacteria as a Regulator of Metabolic Enzyme Complexes. mBio 8, e01096–17 (2017).

14. Li, H. et al. Heat and Pressure Resistance in Escherichia coli Relates to Protein Folding and Aggregation. Front. Microbiol. 11, 111 (2020).

15. Lin, M.-H. et al. A New Tool to Reveal Bacterial Signaling Mechanisms in Antibiotic Treatment and Resistance. Mol. Cell. Proteomics 17, 2496–2507 (2018).

16. Hakobyan, A., Schneider, M. B., Liesack, W. & Glatter, T. Efficient Tandem LysC/Trypsin Digestion in Detergent Conditions. PROTEOMICS 19, 1900136 (2019).

17. Makukhin, N. et al. Resolving oxidative damage to methionine by an unexpected membrane-associated stereoselective reductase discovered using chiral fluorescent probes. FEBS J. 286, 4024–4035 (2019).

18. Couté, Y., Bruley, C. & Burger, T. Beyond Target–Decoy Competition: Stable Validation of Peptide and Protein Identifications in Mass Spectrometry-Based Discovery Proteomics. Anal. Chem. 92, 14898–14906 (2020).

19. Lima, D. B. et al. ProteoCombiner: integrating bottom-up with top-down proteomics data for improved proteoform assessment. Bioinformatics 37, 2206–2208 (2021).

20. Fortuin, S., Iradukunda, J., Nel, A. J., Blackburn, J. M. & Soares, N. C. Liquid chromatography mass spectrometry-based proteomics of Escherichia coli single colony. MethodsX 8, 101277 (2021).

21. Zuily, L. et al. Copper Induces Protein Aggregation, a Toxic Process Compensated by Molecular Chaperones. mBio 13, e03251–21 (2022).

22. Jiang, M. et al. Reductions in bacterial viability stimulate the production of Extra-intestinal Pathogenic Escherichia coli (ExPEC) cytoplasm-carrying Extracellular Vesicles (EVs). PLoS Pathog. 18, e1010908 (2022).

23. Barraud, N. et al. Lifestyle-specific S-nitrosylation of protein cysteine thiols regulates Escherichia coli biofilm formation and resistance to oxidative stress. Npj Biofilms Microbiomes 7, 34 (2021).

24. Donati, S. et al. Multi-omics Analysis of CRISPRi-Knockdowns Identifies Mechanisms that Buffer Decreases of Enzymes in E. coli Metabolism. Cell Syst. 12, 56–67.e6 (2021).

25. Seeger, C. et al. The Subcellular Proteome of a Planctomycetes Bacterium Shows That Newly Evolved Proteins Have Distinct Fractionation Patterns. Front. Microbiol. 12, 643045 (2021).

26. Moyer, T. B., Purvis, A. L., Wommack, A. J. & Hicks, L. M. Proteomic response of Escherichia coli to a membrane lytic and iron chelating truncated Amaranthus tricolor defensin. BMC Microbiol. 21, 110 (2021).

27. Sadecki, P. W. et al. Evolution of Polymyxin Resistance Regulates Colibactin Production in *Escherichia coli*. ACS Chem. Biol. 16, 1243–1254 (2021).

28. Ruan, X. et al. Effect of resveratrol on the biofilm formation and physiological properties of avian pathogenic Escherichia coli. J. Proteomics 249, 104357 (2021).

29. Li, Z. et al. Biodistribution of 89Zr-DFO-labeled avian pathogenic Escherichia coli outer membrane vesicles by PET imaging in chickens. Poult. Sci. 102, 102364 (2023).

30. Kasari, M., Kasari, V., Kärmas, M. & Jõers, A. Decoupling Growth and Production by Removing the Origin of Replication from a Bacterial Chromosome. ACS Synth. Biol. 11, 2610–2622 (2022).

31. Dong, H. et al. YiaC and CobB regulate lysine lactylation in Escherichia coli. Nat. Commun. 13, 6628 (2022).

32. Feng, J. et al. Proteomics Reveal the Effect of Exogenous Electrons on Electroactive Escherichia coli. Front. Microbiol. 13, 815366 (2022).

33. Chen, W. et al. Dimethyl phthalate inhibits the growth of Escherichia coli K-12 by regulating sugar transport and energy metabolism. Environ. Sci. Pollut. Res. Int. 30, 13702–13710 (2023).

34. Seo, H., Giannone, R. J., Yang, Y.-H. & Trinh, C. T. Proteome reallocation enables the selective de novo biosynthesis of non-linear, branched-chain acetate esters. Metab. Eng. 73, 38–49 (2022).

35. Fages-Lartaud, M., Tietze, L., Elie, F., Lale, R. & Hohmann-Marriott, M. F. mCherry contains a fluorescent protein isoform that interferes with its reporter function. Front. Bioeng. Biotechnol. 10, 892138 (2022).

36. Disela, R., Bussy, O. L., Geldhof, G., Pabst, M. & Ottens, M. Characterisation of the E. coli HMS174 and BLR host cell proteome to guide purification process development. Biotechnol. J. 18, e2300068 (2023).

37. Young, J. W., Zhao, Z., Wason, I. S. & Duong van Hoa, F. A Dual Detergent Strategy to Capture a Bacterial Outer Membrane Proteome in Peptidiscs for Characterization by Mass Spectrometry and Binding Assays. J. Proteome Res. 22, 1537–1545 (2023).

38. Shi, H. et al. CycA-Dependent Glycine Assimilation Is Connected to Novobiocin Susceptibility in Escherichia coli. Microbiol. Spectr. 10, e0250122 (2022).

39. Marshall, S. A. et al. The broccoli-derived antioxidant sulforaphane changes the growth of gastrointestinal microbiota, allowing for the production of anti-inflammatory metabolites. J. Funct. Foods 107, 105645 (2023).

40. Yu, M. S. C. et al. The proteome of bacterial membrane vesicles in Escherichia coli—a time course comparison study in two different media. Front. Microbiol. 15, 1361270 (2024).

41. Végvári, Á., Zhang, X. & Zubarev, R. A. Toward Single Bacterium Proteomics. J. Am. Soc. Mass Spectrom. 34, 2098–2106 (2023).

42. He, Y., Jin, H. & Ju, F. Toxicological effects and underlying mechanisms of chlorination-derived metformin byproducts in Escherichia coli. Sci. Total Environ. 905, 167281 (2023).

43. Recacha, E. et al. Impact of suppression of the SOS response on protein expression in clinical isolates of Escherichia coli under antimicrobial pressure of ciprofloxacin. Front. Microbiol. 15, 1379534 (2024).

44. Claeys, T. et al. lesSDRF is more: maximizing the value of proteomics data through streamlined metadata annotation. Nat. Commun. 14, 6743 (2023).

45. Dai, C. et al. A proteomics sample metadata representation for multiomics integration and big data analysis. Nat. Commun. 12, 5854 (2021).

46. Frankenfield, A. M., Ni, J., Ahmed, M. & Hao, L. Protein Contaminants Matter: Building Universal Protein Contaminant Libraries for DDA and DIA Proteomics. J. Proteome Res. 21, 2104–2113 (2022).

47. Kong, A. T., Leprevost, F. V., Avtonomov, D. M., Mellacheruvu, D. & Nesvizhskii, A. I. MSFragger: ultrafast and comprehensive peptide identification in mass spectrometry–based proteomics. Nat. Methods 14, 513–520 (2017).

48. Martens, L. et al. mzML—a Community Standard for Mass Spectrometry Data. Mol. Cell. Proteomics 10, R110.000133 (2011).

49. Hulstaert, N. et al. ThermoRawFileParser: Modular, Scalable, and Cross-Platform RAW File Conversion. J. Proteome Res. 19, 537–542 (2020).

50. Adusumilli, R. & Mallick, P. Data Conversion with ProteoWizard msConvert. in Proteomics (eds Comai, L., Katz, J. E. & Mallick, P.) vol. 1550 339–368 (Springer New York, New York, NY, 2017).

51. Käll, L., Canterbury, J. D., Weston, J., Noble, W. S. & MacCoss, M. J. Semi-supervised learning for peptide identification from shotgun proteomics datasets. Nat. Methods 4, 923–925 (2007).

52. Beausoleil, S. A., Villén, J., Gerber, S. A., Rush, J. & Gygi, S. P. A probability-based approach for high-throughput protein phosphorylation analysis and site localization. Nat. Biotechnol. 24, 1285–1292 (2006).

53. Barente, A. S. & Villén, J. A Python Package for the Localization of Protein Modifications in Mass Spectrometry Data. J. Proteome Res. 22, 501–507 (2023).

54. Reiter, L. et al. Protein Identification False Discovery Rates for Very Large Proteomics Data Sets Generated by Tandem Mass Spectrometry. Mol. Cell. Proteomics 8, 2405–2417 (2009).

55. Keller, A., Nesvizhskii, A. I., Kolker, E. & Aebersold, R. Empirical Statistical Model To Estimate the Accuracy of Peptide Identifications Made by MS/MS and Database Search. Anal. Chem. 74, 5383–5392 (2002).

56. Krug, K. et al. Deep Coverage of the Escherichia coli Proteome Enables the Assessment of False Discovery Rates in Simple Proteogenomic Experiments. Mol. Cell. Proteomics 12, 3420–3430 (2013).

57. van Wijk, K. J. et al. The Arabidopsis PeptideAtlas: Harnessing worldwide proteomics data to create a comprehensive community proteomics resource. Plant Cell 33, 3421–3453 (2021).

58. Broadbent, J. A., Broszczak, D. A., Tennakoon, I. U. K. & Huygens, F. Pan-proteomics, a concept for unifying quantitative proteome measurements when comparing closely-related bacterial strains. Expert Rev. Proteomics 13, 355–365 (2016).

59. Yu, N. Y. et al. PSORTb 3.0: improved protein subcellular localization prediction with refined localization subcategories and predictive capabilities for all prokaryotes. Bioinformatics 26, 1608–1615 (2010).

60. Levitsky, L. I., Klein, J. A., Ivanov, M. V. & Gorshkov, M. V. Pyteomics 4.0: Five Years of Development of a Python Proteomics Framework. J. Proteome Res. 18, 709–714 (2019).

61. Lin, Z. et al. Evolutionary-scale prediction of atomic level protein structure with a language model. Preprint at 10.1101/2022.07.20.500902 (2022).

62. Kanehisa, M. KEGG: Kyoto Encyclopedia of Genes and Genomes. Nucleic Acids Res. 28, 27–30 (2000).

63. Maglott, D. Entrez Gene: gene-centered information at NCBI. Nucleic Acids Res. 33, D54–D58 (2004).

64. Hackl, S. et al. PTMVision: An Interactive Visualization Webserver for Post-translational Modifications of Proteins. J. Proteome Res. 24, 919–928 (2025).

65. Łos, M. et al. Bacteriophage contamination: is there a simple method to reduce its deleterious effects in laboratory cultures and biotechnological factories? J. Appl. Genet. 45, 111–120 (2004).

66. Chen, J., Li, Y., Zhang, K. & Wang, H. Whole-Genome Sequence of Phage-Resistant Strain Escherichia coli DH5α. Genome Announc. 6, e00097–18 (2018).

67. Rostøl, J. T. & Marraffini, L. (Ph)ighting Phages: How Bacteria Resist Their Parasites. Cell Host Microbe 25, 184–194 (2019).

68. Camprubí-Font, C. & Martinez-Medina, M. Why the discovery of adherent-invasive *Escherichia coli* molecular markers is so challenging? World J. Biol. Chem. 11, 1–13 (2020).

69. Macek, B. et al. Protein post-translational modifications in bacteria. Nat. Rev. Microbiol. 17, 651–664 (2019).

70. Creasy, D. M. & Cottrell, J. S. Unimod: Protein modifications for mass spectrometry. PROTEOMICS 4, 1534–1536 (2004).

71. Bludau, I. et al. The structural context of posttranslational modifications at a proteome-wide scale. PLOS Biol. 20, e3001636 (2022).

72. Ramasamy, P., Vandermarliere, E., Vranken, W. F. & Martens, L. Panoramic Perspective on Human Phosphosites. J. Proteome Res. 21, 1894–1915 (2022).

73. Helbig, K., Bleuel, C., Krauss, G. J. & Nies, D. H. Glutathione and Transition-Metal Homeostasis in *Escherichia coli*. J. Bacteriol. 190, 5431–5438 (2008).

74. Macek, B. et al. Protein post-translational modifications in bacteria. Nat. Rev. Microbiol. 17, 651–664 (2019).

75. Perez-Riverol, Y. et al. The PRIDE database at 20 years: 2025 update. Nucleic Acids Res. 53, D543–D553 (2025).

